# BR-bodies provide selectively permeable condensates that stimulate mRNA decay and prevent release of decay intermediates

**DOI:** 10.1101/690628

**Authors:** Nadra Al-Husini, Dylan T. Tomares, Zechariah Pfaffenberger, Nisansala S. Muthunayake, Mohammad A. Samad, Tiancheng Zuo, Obaidah Bitar, James R. Aretakis, Mohammed-Husain M. Bharmal, Alisa Gega, Julie S. Biteen, W. Seth Childers, Jared M. Schrader

## Abstract

Biomolecular condensates play a key role in organizing RNAs and proteins into membraneless organelles. Bacterial RNP-bodies (BR-bodies) are a biomolecular condensate containing the RNA degradosome mRNA decay machinery, but the biochemical function of such organization remains poorly defined. Here we define the RNA substrates of BR-bodies through enrichment of the bodies followed by RNA-seq. We find that long, poorly translated mRNAs, small RNAs, and antisense RNAs are the main substrates, while rRNA, tRNA, and other conserved ncRNAs are excluded from these bodies. BR-bodies stimulate the mRNA decay rate of enriched mRNAs, helping to reshape the cellular mRNA pool. We also observe that BR-body formation promotes complete mRNA decay, avoiding the build-up of toxic endo-cleaved mRNA decay intermediates. The combined selective permeability of BR-bodies for both, enzymes and substrates together with the stimulation of the sub-steps of mRNA decay provide an effective organization strategy for bacterial mRNA decay.

## Introduction

The bacterial cytoplasm has been thought to be poorly organized due to the broad lack of membrane bound organelles which compartmentalize biochemical pathways by selective membrane permeability (Abbondanzieri and Meyer, 2019; Kerfeld et al., 2018; Surovtsev and Jacobs-Wagner, 2018). Rapid advances have recently demonstrated biomolecular condensation promotes the formation of membraneless compartments that selectively concentrate and organize proteins and nucleic acids (Banani et al., 2017; Courchaine et al., 2016; Shin and Brangwynne, 2017). Biomolecular condensation occurs through liquid-liquid phase separation of intrinsically disordered scaffolding proteins together with RNA resulting in cytosolic liquid-like droplets. The formation of these droplets is driven by networks of weak multivalent interactions between scaffolding proteins and their recruited client proteins (Banani et al., 2017). BR-bodies are the first biomolecular condensates in bacteria that were shown to directly assemble through liquid-liquid phase separation(Al-Husini et al., 2018). Phase separation occurs through the RNase E scaffold protein’s intrinsically disordered region (IDR) together with RNA degradosome client proteins and RNA (Al-Husini et al., 2018). These BR-body forming capabilities were previously shown to increase cell survival during stress exposure (Al-Husini et al., 2018), however, the biochemical function of these BR-bodies in organizing mRNA decay is poorly, understood.

Many eukaryotic biomolecular condensates have been shown to concentrate a subset of cellular proteins and RNAs, such as nucleoli with rRNA transcription and processing machinery, cajal bodies with snRNP assembly, and germ granules with germ cell-related mRNPs (Banani et al., 2017; Courchaine et al., 2016; Shin and Brangwynne, 2017; Weber, 2017). Most similar to BR-bodies, eukaryotic P-bodies organize the mRNA decay machinery by selective permeability to decay related proteins including decapping enzymes, deadenylation factors, translation repressors, and the major cytoplasmic nuclease Xrn1 (Hubstenberger et al., 2017). Stress granules also share similarity with BR-bodies: these granules are assembled with Xrn1 and some overlapping translation repressors, but they lack the decapping factors and deadenylases and instead contain translation initiation factors (Jain et al., 2016). Recently it, was revealed in purified stress granule cores that many long, poorly translated mRNAs are enriched in stress granules, showing that stress granules also exhibit selective permeability for substrate RNAs (Khong et al., 2017). Interestingly, this selectivity may come from the RNA itself and its abilities to form intermolecular base pairs, yielding toxic RNA assemblies (Van Treeck et al., 2018). While both P-bodies and stress granules share poorly translated mRNAs as substrates, P-bodies tend to be devoid of all ribosomes (Hubstenberger et al., 2017), whereas stress granules contain stalled initiation complexes but lack the large ribosome subunit (Reineke et al., 2012). Additionally, translational repressors such as microRNAs (miRNAs) associate in P-bodies (Eulalio et al., 2008), yet the interplay with both ribosomes or, inhibitory bacterial small RNAs (sRNAs) which also silence mRNAs through imperfect base-pairing is, unknown for BR-bodies.

The biochemical function of P-bodies and stress granules on mRNA decay is less well, understood. P-bodies have been found to stimulate mRNA decay (Sheth and Parker, 2003) and recent, mathematical models have found that smaller P-bodies might act as more efficient sites of mRNA decay, (Pitchiaya et al., 2019), yet P-bodies are not necessary for mRNA decay (Eulalio et al., 2007). P-bodies, can also store untranslated mRNAs (Hubstenberger et al., 2017; Sheth and Parker, 2003), suggesting, that the mRNA fate may depend on the specific mRNA identity and context of cellular conditions (Wang et al., 2018). How stress granules affect mRNA decay and or mRNA function remains to be established,, but their lack of deadenylases and decapping factors suggests that they are more likely utilized for, mRNA storage than for decay (Ivanov et al., 2019; Protter and Parker, 2016).

In most bacterial mRNAs, decay initiates by endonuclease cleavage by RNase E, which provides, the rate-limiting step of the process (Bandyra and Luisi, 2018; Hui et al., 2014; Mohanty and Kushner, 2016). Upon endonuclease cleavage, 3’-5’ exoribonucleases degrade the mRNA decay intermediates, generated by RNase E cleavage (Bandyra and Luisi, 2018; Hui et al., 2014; Mohanty and Kushner, 2016). RNase E’s scaffolding activity of the RNA degradosome proteins, including conserved 3’-5’, exoribonuclease PNPase, help to stimulate the decay process (Ait-Bara and Carpousis, 2015; Bandyra, and Luisi, 2018; Lopez et al., 1999). RNase E and 3’-5’ exonucleases work cooperatively with DEAD-box, RNA helicases to ensure mRNA decay intermediates do not accumulate, which are not detectable in wild, type cells and can only be detected when the IDR of RNase E or exoribonucleases are mutated (Coburn et al., 1999; Khemici and Carpousis, 2004; Morita et al., 2004). *Caulobacter crescentus* RNase E is known to recruit PNPase, RhlB, and RNase D into its RNA degradosome and BR-bodies (Al-Husini et al., 2018; Hardwick et al., 2011; Voss et al., 2014). The assembly of BR-bodies independent of RNA degradosome protein association was found to stimulate mRNA decay for the RNase E mRNA (Al-Husini et al., 2018), however, the exact mRNA decay steps stimulated by biomolecular condensation are not understood. Additionally, RNase E is known to control global mRNA decay in *E. coli* (Clarke et al., 2014; Hammarlof et al., 2015; Ono and Kuwano, 1979), with mutations blocking foci-formation leading to a slowdown in global decay rates (Hadjeras et al., 2019; Lopez et al., 1999), yet the entire collection of RNA substrates that BR-bodies act on in *C. crescentus* has not yet been defined.

In this study, we combine BR-body enrichment with RNA sequencing (RNA-seq) to define the RNA substrates of BR-bodies, and we use global mRNA half-life profiling of BR-body mutant strains to define their biochemical roles in facilitating mRNA decay. BR-body enrichment defines substrates predominantly as mRNAs and small RNAs (sRNAs) that act like miRNAs to silence mRNAs by base pairing. BR-body mRNA substrates are predominantly longer more poorly translated mRNAs, and we find that the RNA length is an intrinsic property that stimulates BR-body assembly *in vitro*. We find rRNA and the nucleoid are physically excluded from BR-bodies, thereby providing selective permeability of target substrates and rejecting molecules that compete for mRNA substrates. Additionally, global mRNA half-life profiling reveals that BR-bodies stimulate decay of BR-body enriched mRNAs by stimulating both the initial endo-cleavage by RNase E and the subsequent exonucleolytic decay step by degradosome associated exonucleases. This study therefore provides new functional insights into how the organization of the mRNA decay machinery into biomolecular condensates can facilitate robust biochemical pathways.

## Results

### RNase E cleavage stimulates rapid mRNA decay in *C. crescentus*

As *E. coli* RNase E provides the rate limiting cleavage in mRNA decay (Hammarlof et al., 2015; Ono and Kuwano, 1979), we sought to determine if the *C. crescentus* enzyme also provides this critical function. We measured and compared the mRNA half-lives using a modified RNA-seq assay for the wild type RNase E and a strain with an RNase E active site mutation (ASM) during exponential growth (Callaghan et al., 2005) (Fig 1A). The resulting bulk mRNA half-lives were 3.6 min for the wild type, and 7.6 min for the ASM, suggesting that endonuclease activity stimulates bulk mRNA decay (Fig S1A). The approximately 2-fold slowdown in mRNA decay rate observed with the ASM is slightly lower than the 5-fold decrease observed in an *E. coli* RNase ETS strain (Ono and Kuwano, 1979) perhaps due to the ASM mutant’s ability to maintain degradosome formation. While bulk estimates of sRNAs and tRNAs remain stable, we observed a 3.0 min bulk half-life for cis-encoded antisense RNAs (asRNAs), whose lifetime increased slightly to 5.0 minutes in the ASM mutant. By examining the individual mRNA decay rates, we found the median mRNA decay rate is 1.6 minutes with the active RNase E, while the median mRNA decay rate rose to 3.7 minutes in the ASM mutant (Fig 1B). The median half-life is likely overestimated in the wild-type because we measure 553 more mRNA half-lives in the ASM strain and these mRNAs were predominantly undetectable by the wild type’s 3 min time-point. Across individual mRNAs we find that 96% of mRNAs have a longer decay rate in the ASM strain as compared to wild type with an average slowdown of 2.4 fold (Fig S1), showing that entry into mRNA decay by RNase E also stimulates global mRNA decay in *C. crescentus* (Fig 1B, S1B). ASM mutation did not affect median RNA half-lives across sRNAs or asRNAs, suggesting that these RNAs may be degraded by other RNases (Fig 1B).

**Figure 1.**
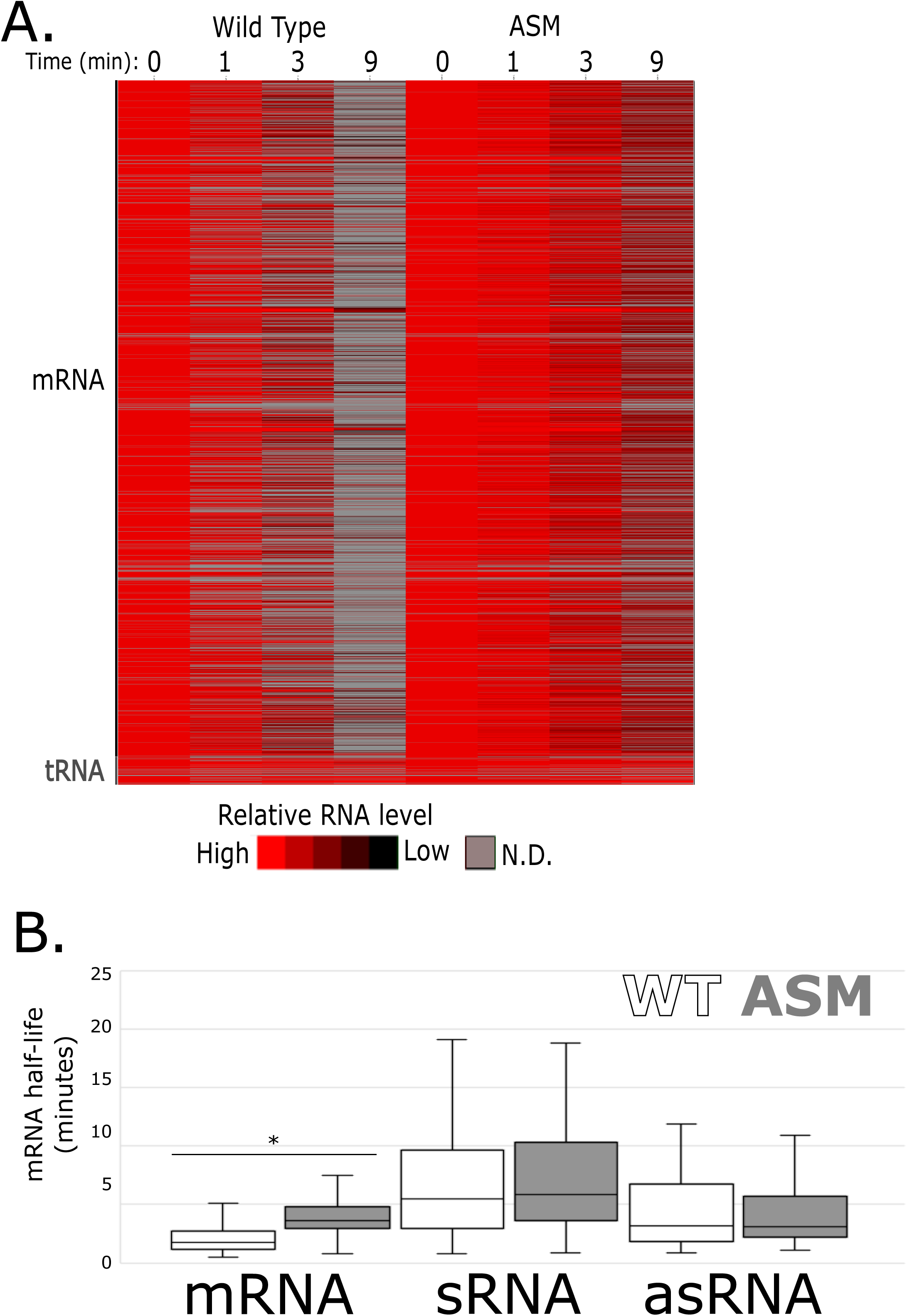
RNase E cleavage is needed for rapid mRNA decay in *C. crescentus*. A.) RNA-seq measurement of wild type (WT, JS38) or active site mutant RNase E variant (ASM, JS299) after treatment with 200 µg/mL rifampicin for the indicated amount of time. Each row represents a different transcript whose RNA level is normalized to the level in untreated cells. Grey color represents a value too low to be determined. B.) Box-plots of RNA half-lives based on the data in panel A. Two-tailed T-test between FL and ASM mRNA decay rates with uneven variance yield a p-value of 1.3 × 10^-23^.

### BR-bodies engage on longer poorly translated mRNAs

As RNase E’s endonuclease activity stimulates bulk mRNA-decay, we sought to determine the cellular RNA substrates of BR-bodies. Importantly, as native BR-bodies are highly labile and not detectible in a cell lysate, we utilized the ASM mutant of RNase E which blocks dissolution of BR-bodies and allowed them to be stabilized during enrichment (Al-Husini et al., 2018). Initial affinity purification attempts of BR-bodies by an N-terminal HA-tagged ASM yielded highly purified RNase E, however, the bodies dissolved during the hours-long incubation times of purification and RNA was not detectable (Fig S2). Therefore we performed an enrichment of BR-bodies by rapidly separating BR-bodies away from cellular contents by differential centrifugation, similar to the procedure used to isolate stress granule cores (Khong et al., 2018; Wheeler et al., 2016) (Fig S2). Enriched BR-body RNA levels were then compared to RNA levels in the cell lysate using RNA-seq (Fig 2). By examining the fraction of reads among mRNAs and non-coding RNAs we noticed that the lysate contained 69.7% ncRNAs (63.4% rRNA, 5.8% tRNA, 0.005% sRNA, and 0.002% asRNA) and 25.7% mRNAs, while the BR-body enriched samples contained 13.3% ncRNAs (12.7% rRNA, 0.005% tRNA, 0.007% sRNA, and 0.006% asRNA) and 63.4% mRNA (Fig 2C).

**Figure 2.**
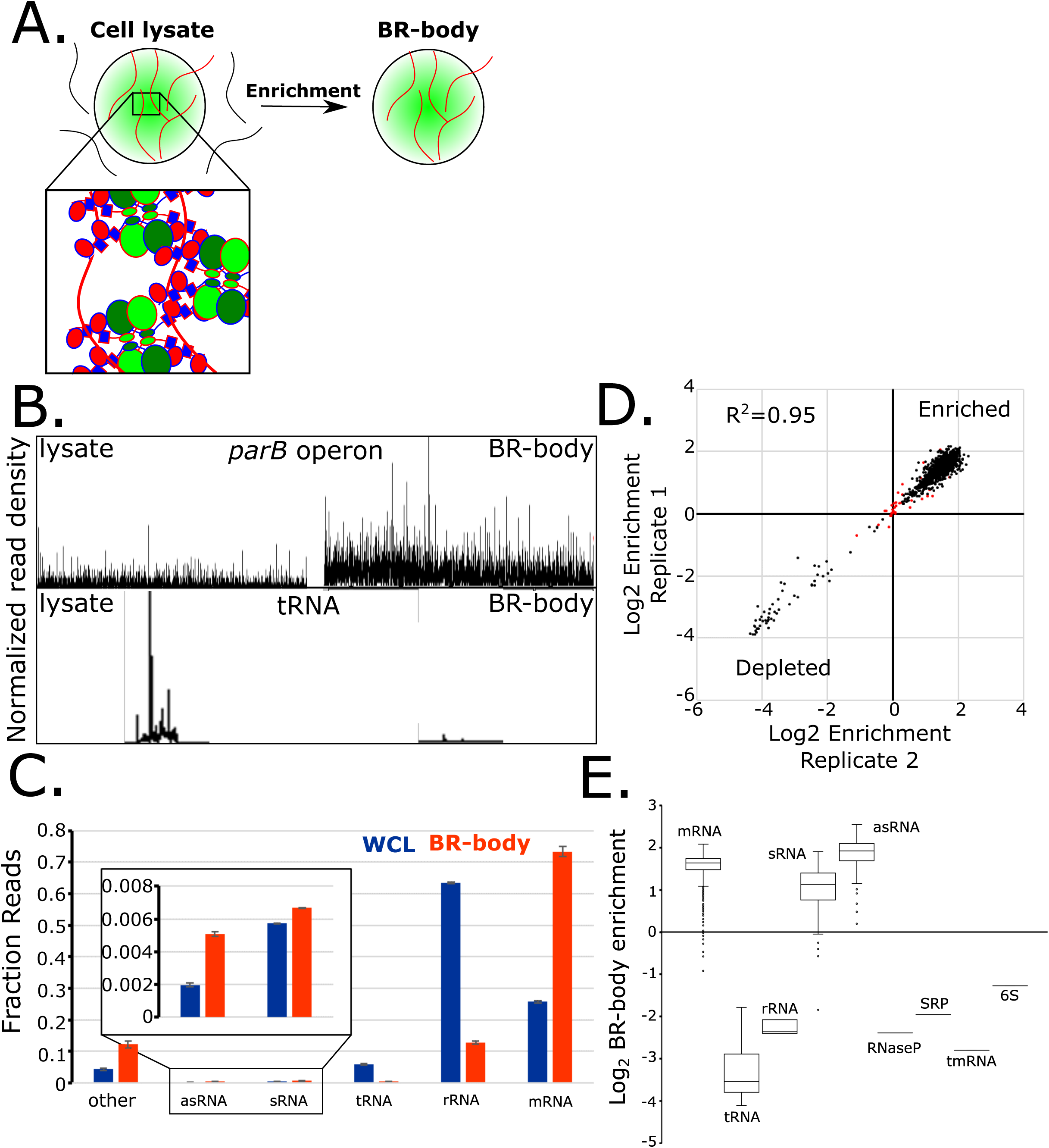
BR-bodies are enriched for mRNAs. A.) Differential centrifugation-based enrichment of BR-bodies. An aliquot of the cell lysate and enriched BR-bodies were RNA-extracted and libraries for RNA-seq were generated and sequenced. B.) Normalized read density for the *parB* operon (top) and for a tRNA gene (bottom). C.) Fraction of RNA-seq reads mapping to each RNA category from whole-cell lysate (WCL, blue) or BR-body enriched samples (orange). D.) Log2 ratio of the BR-body enriched sample RPKM compared to the cell lysate RPKM values of two biological replicates. RNAs with p-values >0.05 are colored in red Fraction of total reads mapping to non-coding RNA (blue) and mRNA (orange) in the lysate and in the enriched BR-body samples. E.) Box plots of BR-body enrichment for RNAs of each RNA category.

To explore which RNA species were enriched or depleted, we compared the log_2_ ratio of enrichment (BR-body read density / total-RNA read density) for each RNA (Fig 2D, Table S1). Two biological replicates of this assay gave relatively similar agreement in the level of BR-body enrichment observed with R^2^ = 0.95 (Fig 2D). We noticed that many conserved ncRNAs, including tRNAs (−3.5 median enrichment), rRNA (−2.3), RNase P (−2.4), tmRNA (−2.8), 6S RNA (−1.3), and SRP RNA (−1.9) were highly depleted in BR-bodies (Fig 2E). Conversely, mRNAs (1.6 median enrichment), sRNAs (1.1), and asRNAs (1.9) were predominantly enriched in BR-bodies (Fig 2E). By comparing mRNAs, sRNAs, and asRNAs we noticed that the level of enrichment positively correlated with the length of the RNAs with the strongest effect observed for mRNAs (Fig 3A, 3B). While hundreds of sRNAs are known to exist in *C. crescentus* (Schrader et al., 2014; Zhou et al., 2015), only three have been functionally explored. Two trans-encoded sRNAs who can silence target mRNAs through base pairing were found to be slightly enriched in BR-bodies (*crfA* 0.67 and *chvR* 0.27) (Frohlich et al., 2018; Landt et al., 2010), while another was found to be slightly depleted (*gsrN* -0.24) (Tien et al., 2018). Pairing of sRNAs with mRNAs typically occurs through the Lsm protein Hfq (Santiago-Frangos and Woodson, 2018; Vogel and Luisi, 2011), and we observe that 252/257 of these RNAs that associate with Hfq (Assis et al., 2019) are enriched in BR-bodies (Table S1).

**Figure 3.**
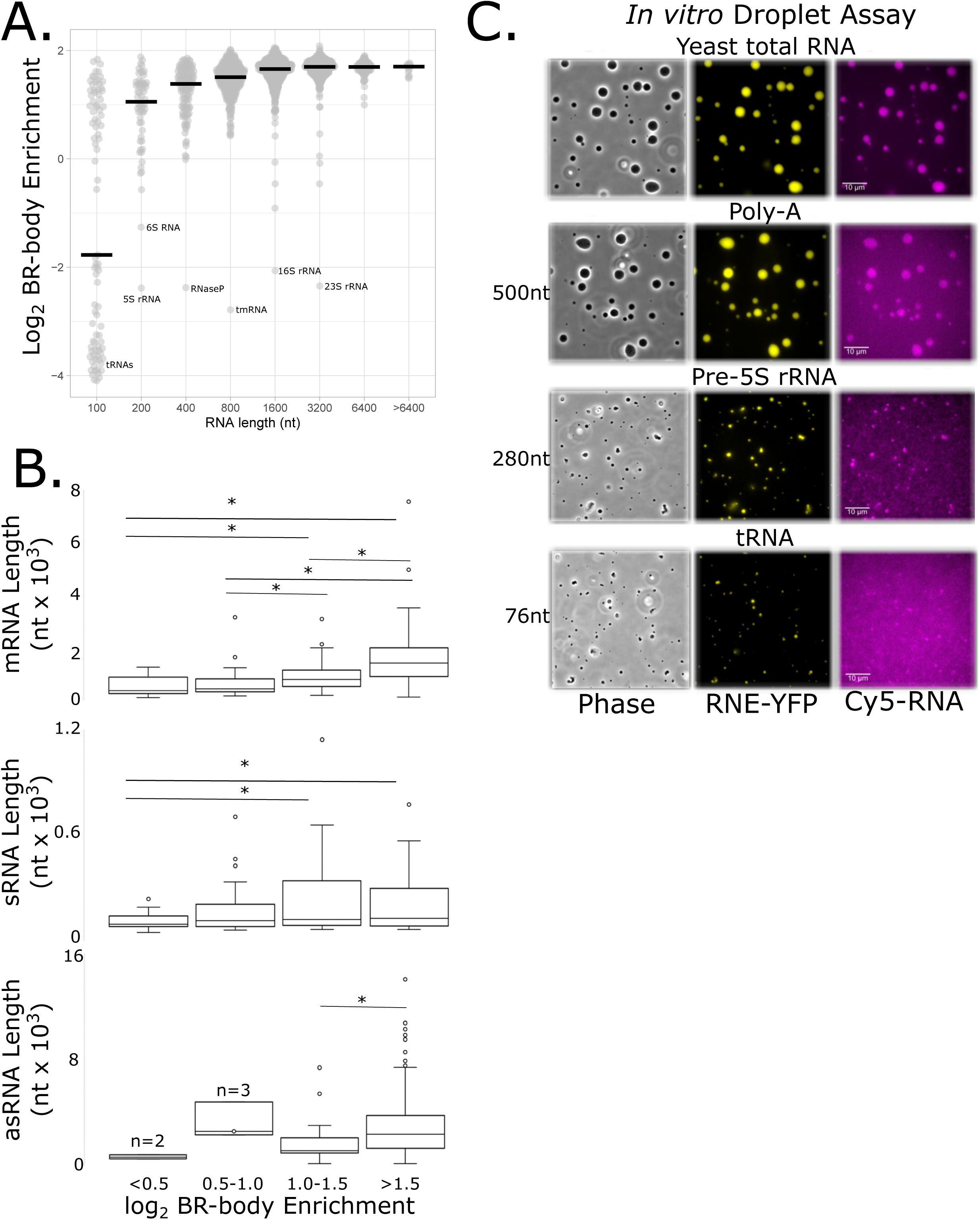
Long mRNAs are preferred substrates of BR-bodies. A.) Log_2_ ratio of the BR-body enriched RPKM vs cell lysate RPKM. RNAs were divided into bins based on their length and the medians of enrichment are highlighted (black bars). Conserved ncRNAs are indicated. B.) Comparison of BR-body enrichment and length for enriched classes of RNAs. Two-tailed T-tests with unequal variances resulting in P-values less than 0.05 are highlighted with asterisks. C.) *In vitro* RNase E CTD-YFP biomolecular condensate assay with the indicated labeled RNA.

As RNase E cleavage stimulates mRNA decay, we explored which mRNA features correlate with BR-body enrichment. The largest correlation coefficients with BR-body enrichment were observed for mRNA abundance (R=-0.51), GC% (R=0.47), mRNA length (R=0.41), translation level (R=-0.40), and codon optimality (nTEI) (R=0.3, Fig S3). The strong negative correlation between mRNA abundance and BR-body enrichment suggests that BR-body mediated decay likely plays a significant role in shaping the cellular mRNA pool. The finding that longer poorly translated mRNAs are highly correlated with BR-body enrichment suggesting a similar mRNA preference as eukaryotic stress granules (Khong et al., 2017; Van Treeck et al., 2018). As the mRNA GC% is also correlated with codon optimality (nTEI) (R=0.57) and sRNAs contain no GC bias in enriched RNAs, this GC% correlation likely relates to the underlying codon usage. Surprisingly, the genes with lower nTEI are poorly enriched in BR-bodies, while those with more highly optimal codons are more highly enriched. Lower correlations were observed for translation efficiency (−0.16), tRNA adaptation index (TAI) (R=0.08), Shine-Dalgarno strength (R=-0.04), and 5’ UTR length (R=0.02) (Fig S3).

While mRNA length correlated rather strongly with BR-body enrichment (R=0.41), we also noted that sRNA and asRNA length correlated positively with BR-enrichment, although to a weaker extent due in part to the limited RNA size ranges of sRNAs and asRNAs (R=0.20 and R=0.22, Fig S4). To explore the intrinsic role of RNA length on BR-body assembly, we added Cy5 dye-labeled RNAs of different sizes to purified condensates formed with the CTD of RNase E which is both necessary and sufficient for BR-body formation (Al-Husini et al., 2018) (Fig 3C). The two samples containing longer Cy5-RNAs (yeast total RNA and poly-A RNA) were efficiently recruited into CTD droplets, while the samples containing shorter cy-5 RNAs (9S pre-rRNA and tRNA) were poorly recruited into the CTD droplets and appeared to partially dissolve them. This enhancement of condensation suggests that the preference of BR-bodies for longer RNA substrates is an intrinsic property that may allow bridging of RNase E proteins to facilitate phase separation.

### BR-bodies are selectively permeable

As BR-bodies compete with ribosomes for free mRNA (Al-Husini et al., 2018), and rRNA was highly depleted from BR-bodies (Fig 2), we explored whether BR-bodies might exclude ribosomes *in vivo*. In *C. crescentus* cells expressing a ribosomal protein L1-eYFP fusion and an RNase E-eCFP fusion as the sole copies, we observe low ribosome density at the sites of BR-bodies (Fig 4A, S5). The occlusion of ribosomes from the BR bodies is further revealed by super-resolution microscopy in fixed *C. crescentus* cells expressing L1-PAmCherry and RNase E-eYFP as the sole copies; these images show that the ribosomes are distributed throughout the cells but are occluded from the RNase E foci. In measurements of 41 cells, we find that only 5% of the L1 localizations occur within 100 nm of an RNase focus (Fig 4B). This anti-localization is contrasted with the control case of cells expressing RNase E-eYFP and the degradosome component aconitase-PAmCherry; as expected, RNase E and aconitase strongly co-localize (Fig S7). As the cytoplasm is filled with the nucleoid in *C. crescentus*, and RNase E was found to associate with the nucleoid (Montero Llopis et al., 2010), we also sought to explore whether the nucleoid was excluded from BR-bodies. We indeed observe occlusion of the nucleoid by DAPI staining (Fig 4C), suggesting BR-bodies create distinct subcellular compartments for mRNA decay (Fig S5). Additionally, by heterologously expressing both the structured catalytic NTD and intrinsically disordered CTD of RNase E in *E. coli* cells, we find that the NTD colocalizes with the *E. coli* nucleoid, while the CTD forms nucleoid occluded condensates throughout the body of the cell (Fig S5).

**Figure 4.**
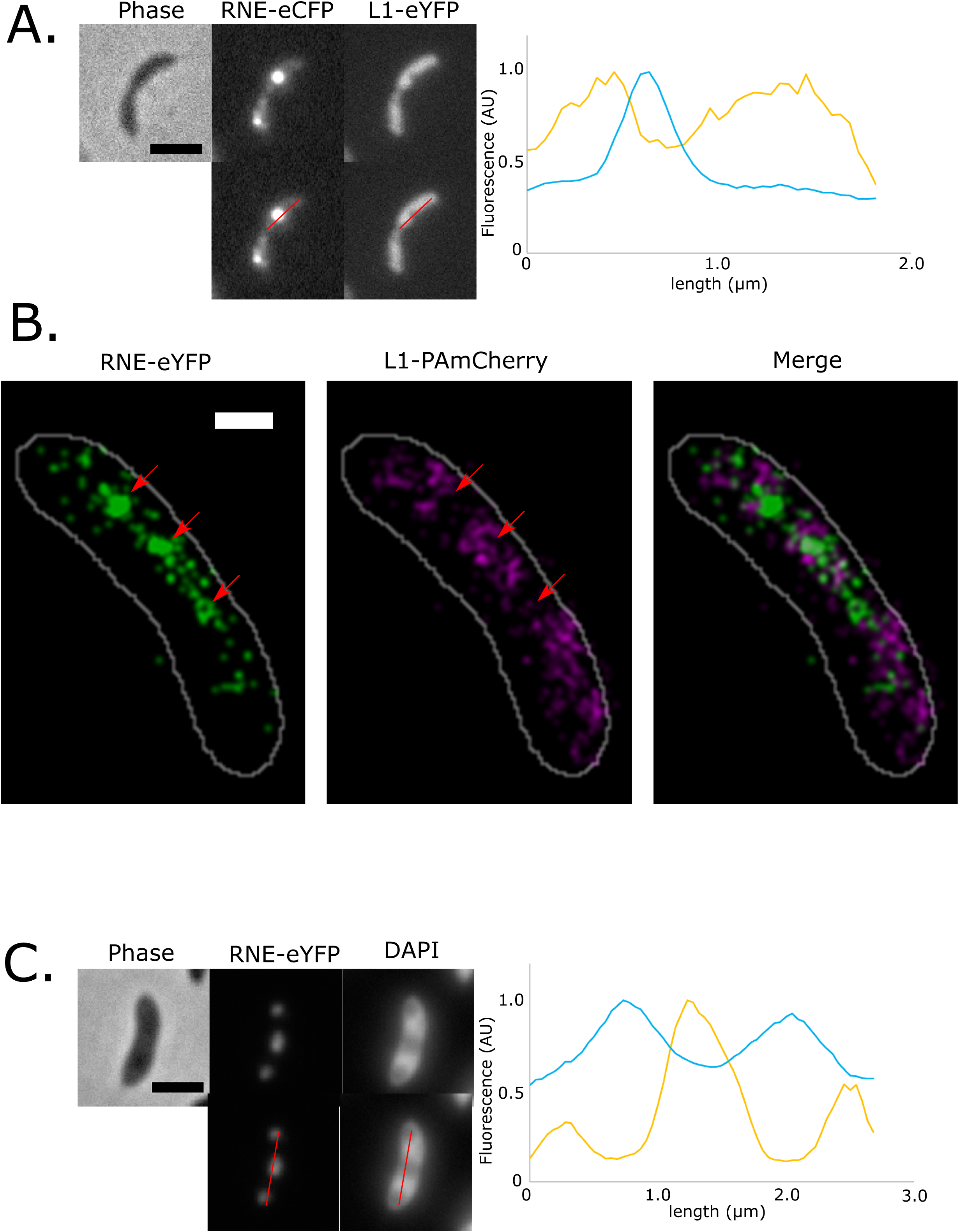
Ribosomes and the nucleoid are occluded from BR-bodies. A.) Dual labeled strain expressing ribosomal protein L1-eYFP and BR-body scaffold RNase E (RNE)-eCFP as the sole copies (JS350). Red line represents position of line trace of signal intensity on right. B.) Super-resolution images of RNE-eYFP (green) and L1-PAmCherry (magenta) (JS545). White cell outlines from phase image. Red arrow indicates position of BR-body. In merged image overlapping eYFP/PAmCherry signal is displayed in white. D.) ASM-YFP (JS299) colocalized with the DAPI stained nucleoid. Scale bars are 2 µm.

We used mRNA FISH and the Ms2 RNA labeling system to colocalize the highly-translated *rsaA* mRNA, which is known to have an unusually long mRNA half-life in *C. crescentus* (*Lau et al*., 2010), together with BR-bodies (Fig 5, S10). By mRNA FISH, we observed that 28% of mRNA foci show some extent of overlap with BR-body foci in diffraction-limited images, however, the *rsaA* mRNA is predominantly observed outside BR-bodies. Visualization of the *rsaA* mRNA by the Ms2-tagged RNA visualization system (Golding and Cox, 2004) using a Ms2 coat protein capsid assembly mutant (LeCuyer et al., 1995) in live cells also showed similar fractions of colocalization with BR-bodies (27%) (Fig 5A). As a negative control, we also colocalized the *rsaA* mRNA with mCherry-PopZ, a polar protein known to occlude ribosomes with no known role in mRNA turnover (Bowman et al., 2008; Ebersbach et al., 2008). Here 12% of *rsaA* mRNA foci colocalized with a mCherry-PopZ foci, suggesting that a significant amount of the *rsaA* mRNA overlap with BR-bodies may be due to the poor resolution obtained by diffraction-limited images relative to the small cell size (Fig 5A). To explain the poor colocalization of BR-bodies with the *rsaA* mRNA (28%) we hypothesized that if mRNA decay occurs within BR-bodies we would expect a low fraction of colocalization due to the rapid internal mRNA decay. Previous work showed that BR-bodies are known to dynamically assemble and disassemble on the sub-minutes scale and that RNA cleavage is needed to disassemble BR-bodies (Al-Husini et al., 2018). Therefore, inhibiting RNase E endonuclease activity may lead to an accumulation of the *rsaA* mRNA in catalytically inactive BR-bodies. In line with this hypothesis, when RNase E’s endonuclease cleavage is blocked, colocalization of the *rsaA* mRNA and BR-bodies increased significantly with a majority of *rsaA* mRNAs (74%) becoming colocalized with BR-bodies (Fig 5B). Taken together, these results suggest that mRNA association with BR-bodies results in a short-lived assembly, wherein mRNA decay is stimulated followed by a rapid disassembly of the BR-body.

**Figure 5.**
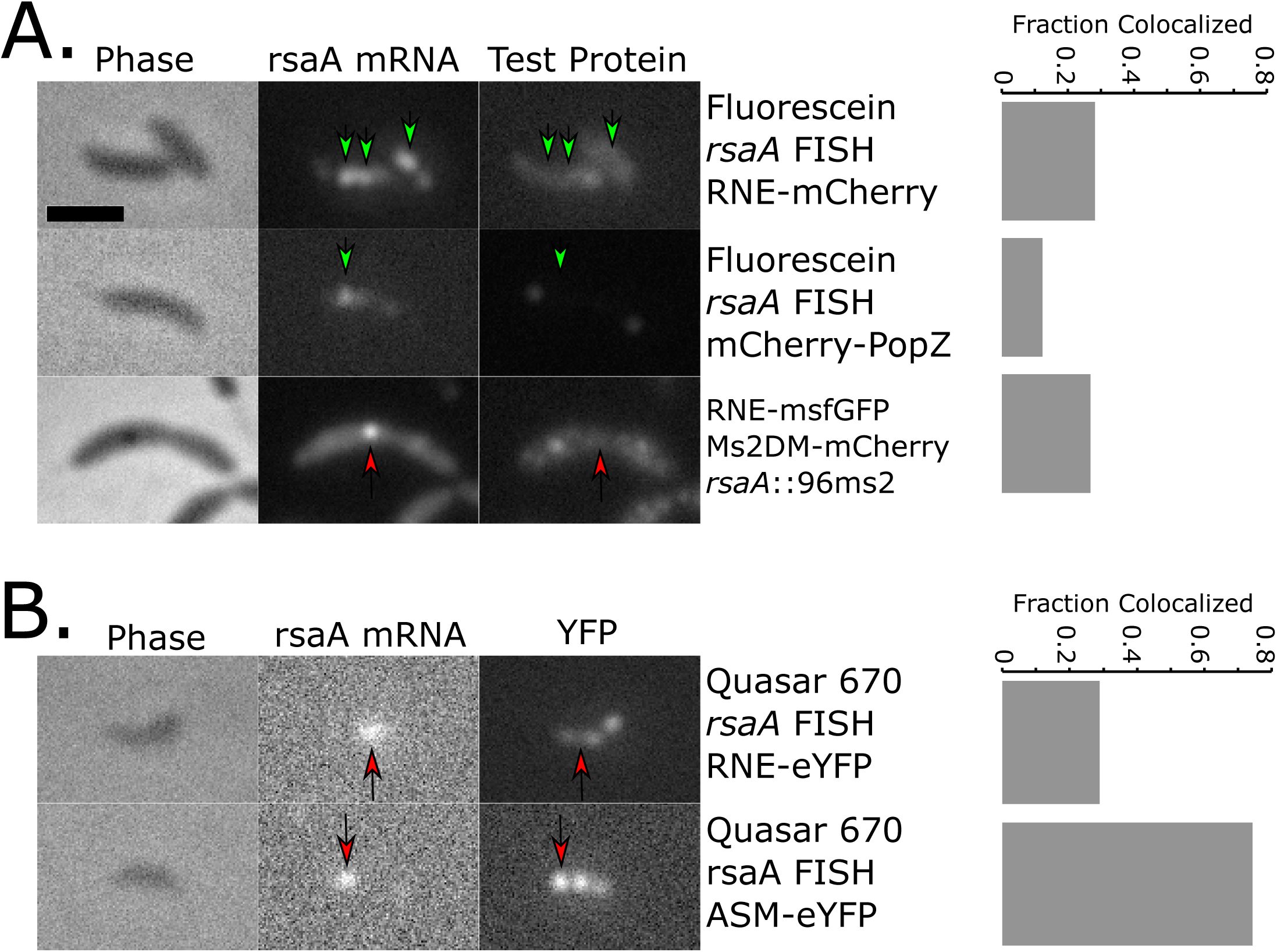
RNase E Endonuclease activity limits *rsaA* mRNA colocalization in BR-bodies. A.) rsaA mRNA weakly colocalizes with BR-bodies. Left, *rsaA* visualization in fixed cells by mRNA FISH or by the Ms2 tagged system in living cells. Fluorescein mRNA FISH probes were probed with either RNE-mCherry (JS403) or mCherry-PopZ (JP369). In live cells the Ms2-coat protein double mutant (Ms2DM)-mCherry fusion with an array of 96 tandem Ms2 RNA hairpins fused to the 3’ end of the rsaA gene was imaged with RNE-msfGFP (JS287). Right, quantitation of the fraction of rsaA mRNA foci colocalized with RNase E or PopZ foci. B.) Left, Quasar 670 mRNA FISH probes (stellaris) were probed with either wild type RNE-eYFP (JS38) or with ASM-eYFP (JS299). Right, quantitation of the fraction of rsaA mRNA foci colocalized with RNase E or ASM foci.

### BR-bodies accelerate RNase E endo cleavage and degradosome exonucleolytic steps

To explore the functional organization of the RNA degradosome into BR-bodies we explored how mutants affecting BR-body assembly modulate mRNA decay using global mRNA half-life profiling (Fig 6). The NTD mutant lacks the IDR and therefore the ability to form a condensate or to scaffold the RNA degradosome. In contrast, the degradosome binding site mutant (ΔDBS) lacks only the scaffolding activity of the IDR for degradosome exoribonucleases, while it retains condensate formation properties allowing the separation of functions (Al-Husini et al., 2018). In the strain expressing wild-type RNase E, the bulk mRNA half-life was the fastest (3.6 min) and mRNAs with lower BR-body enrichment tend to have longer half-lives, suggesting BR-body enrichment stimulates mRNA decay at a global level (Fig 6B).

**Figure 6.**
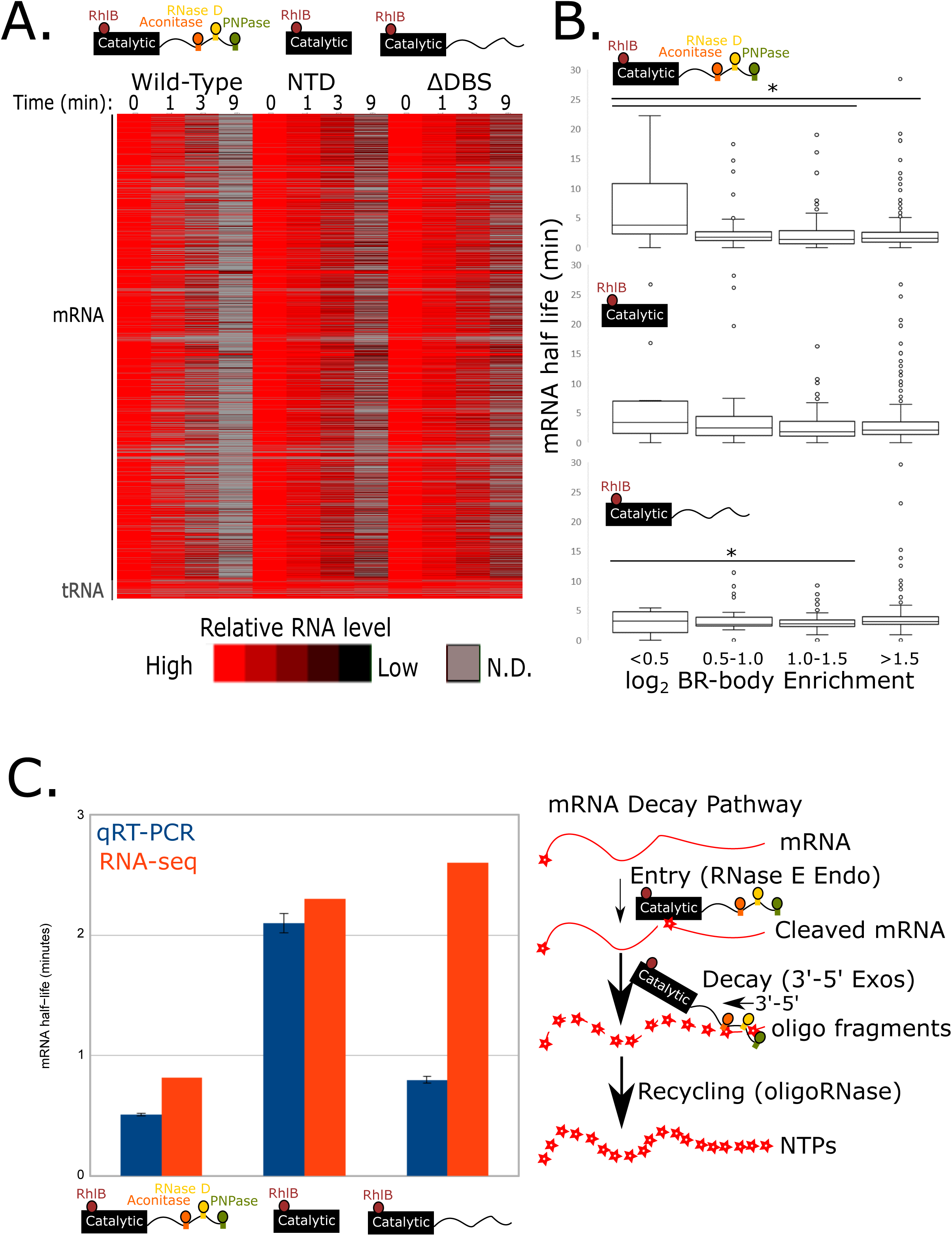
BR-bodies accelerate initial cleavage and exonucleolytic steps of mRNA decay. A.) RNA-seq measurement of wild type (JS38), NTD truncation (JS221), or DBS mutant (JS233) strains after treatment with 200 µg/mL rifampicin for the indicated amount of time. Each row represents a different transcript whose RNA level is normalized to the level in untreated cells. B.) mRNA half-lives for each RNase E mutant across four bins of BR-body enrichment. Only simple mRNAs with a single ORF and TSS are shown. Asterisks indicate samples with p-values ≤0.05. C.) qRT-PCR and RNA-seq half-life measurements for the indicated strains. Each half-life measurement was performed on the same RNA samples split between the two assays (left). Cartoon of the steps of mRNA decay with indicated model of the degradosome (right).

The NTD mutant had a modest bulk slowdown in decay (3.6 to 4.8 min), similar to an *E. coli* RNase E CTD truncation (Lopez et al., 1999). Importantly, the NTD mutant also showed a strong slowdown in half-lives occurring predominantly in mRNAs that are enriched in BR-bodies (Fig 6A,B). The ΔDBS mutant also showed a similar slowdown of bulk mRNA decay (3.6 to 4.5 min) with slower half-lives of mRNAs that are enriched in BR-bodies (Fig 6A,B).

To explore whether different steps in mRNA decay were affected in the RNase E mutants, we used quantitative reverse transcription PCR (qRT-PCR), which examines the integrity of longer pieces of RNA >100nt in length, compared to the shorter 15-50nt fragments measured by RNA-seq. The RNA degradosome facilitates the multi-step mRNA decay process, in which the endonuclease RNase E makes the first initial cleavage followed by exonuclease activity of degradosome associated nucleases (Fig 6C). Therefore, half-lives measured by qRT-PCR will likely be most sensitive to the initial cleavage step, while half-lives measured by RNA-seq will likely be sensitive to both endo-cleavage and the partially cleaved mRNA decay intermediates generated by exoribonucleases (Fig 6C). We then tested the three RNase E variants by RNA-seq and qRT-PCR measurements on the same total RNA samples, yielding dual half-life measurements for 4 substrate mRNAs and the 9S rRNA which is known to be processed by RNase E (Hardwick et al., 2011). For the wild type RNase E, qRT-PCR half-lives ranged from 0.51-0.81 minutes, while RNA-seq measures ranged between 0.71-1.1 minutes. Out of the 4 mRNAs tested, only one showed a half-life that was significantly faster by qRT-PCR than by RNA-seq with a difference of 0.3 minutes (Fig 6C, Table 1). qRT-PCR mRNA half-lives were all slower than wild type for the NTD mutant (ranging from 1.5-2.7 minutes), which were partially restored in the ΔDBS mutant (qRT-PCR mRNA half-lives 0.80-1.9 minutes) suggesting condensation stimulates RNase E endonuclease activity. When comparing to the RNA-seq derived half-lives, 3 out of 4 mRNA half-lives measured for the NTD mutant were found to be significantly faster by qRT-PCR than by RNA-seq with differences on the order of 0.2-0.7 minutes (Fig 6C, Table 1). Finally, for the ΔDBS mutant, 4/4 mRNA half-lives were found to be significantly faster by qRT-PCR than by RNA-seq with larger differences on the order of 1.2-1.8 minutes (Fig 6C, Table 1). Rates of 9S rRNA processing into the 5S rRNA, an essential function of RNase E that does not require the C-terminal IDR (Hardwick et al., 2011), were virtually identical for the wild type and NTD mutants (Table 1), with a significant slowdown in both qRT-PCR and RNA-seq measured in the ΔDBS mutant. The slower mRNA half-lives measured by RNA-seq relative to qRT-PCR for the NTD and ΔDBS mutant suggests that recruitment of degradosome associated exoribonucleases (PNPase and RNase D) into BR-bodies stimulates the secondary exonucleolytic decay steps of endo-cleaved mRNA decay intermediates (Fig 7).

**Figure 7.**
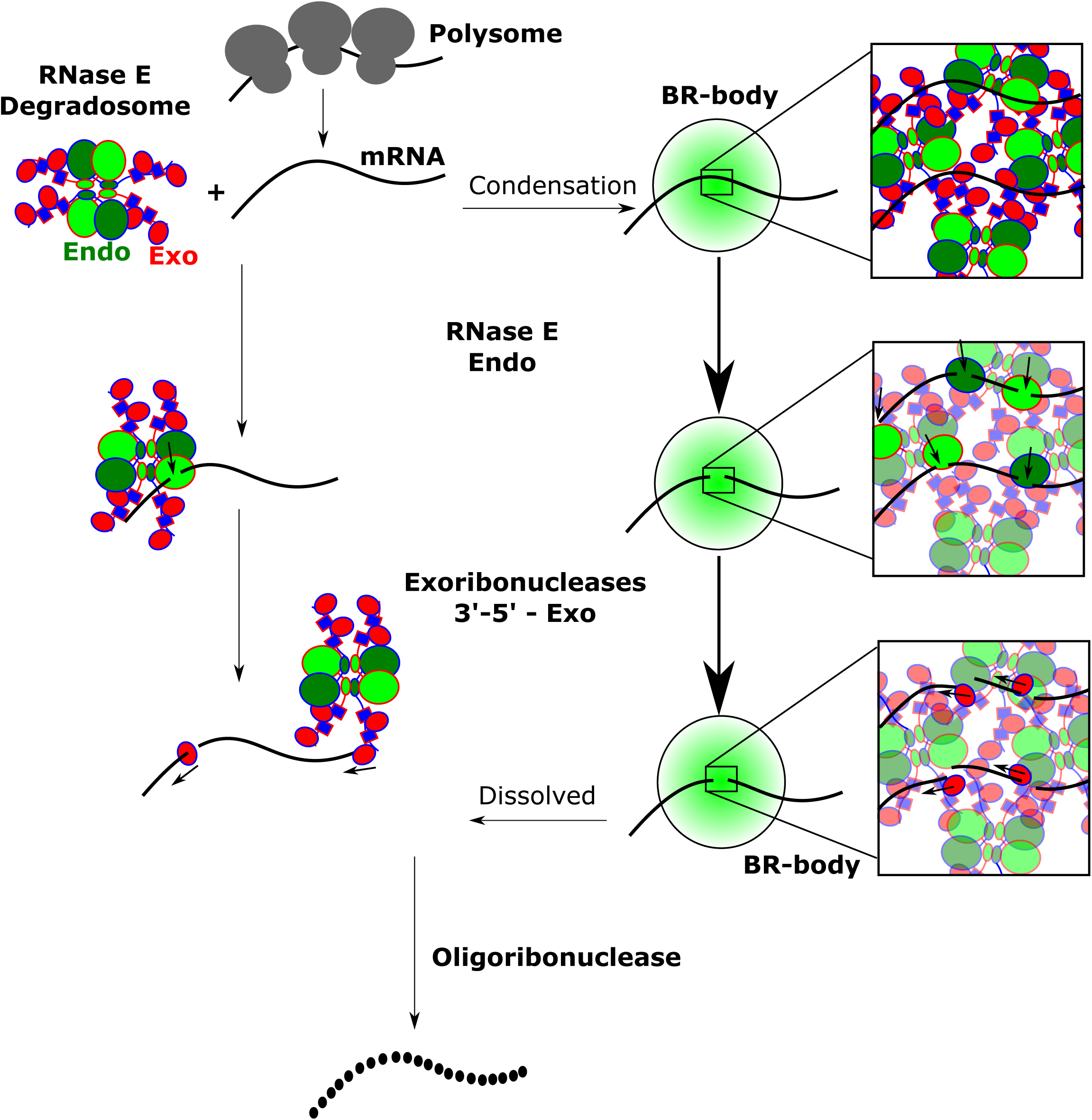
Model of BR-body mediated mRNA decay. As translation levels drop, RNase E can engage on the mRNA to initiate decay. On the left, is the pathway of decay for soluble RNA degradosome where initiation of decay by RNase E and exoribonuclease steps are indicated and can occur slowly. On the right, is the pathway that can occur inside BR-bodies. First, the RNA degradosome and mRNAs condense into a biomolecular condensate. Inside the condensate RNase E endonuclease and degradosome exoribonuclease activity are both accelerated from the high-local concentration. Once the mRNA fragments are cut to a small enough size, the BR-body can dissolve releasing both RNA degradosomes and oligonucleotides that can be turned into nucleotides by oligoribonuclease.

**Table 1.** RNA half-life measurements.

| RNA | RNE<br>variant | RNA half-life (min) | | | $\sigma^{\&}$ | p-value |
| --- | --- | --- | --- | --- | --- | --- |
| | | qRT-PCR | $\sigma$ | RNA-seq | | |
| ***rne***<br>**mRNA** | FL | 0.51* | 0.01 | 0.82 | 0.13 | 3.E-05 |
|  | NTD | 2.1* | 0.14 | 2.3 | 0.13 | 2.E-03 |
| | ΔDBS | 0.80* | 0.05 | 2.6 | 0.58 | 0.E+00 |
| ***gcrA*<sup>#</sup>**<br>**mRNA** | FL | 0.81 | 0.07 | 1.1 | 0.37 | 9.E-02 |
|  | NTD | 2.7 | 0.12 | 3.4 | 0.24 | 4.E-07 |
| | ΔDBS | 1.9 | 0.17 | 3.1 | 0.34 | 3.E-09 |
| ***ctrA***<br>**mRNA** | FL | 0.79 | 0.03 | 0.86 | 0.11 | 1.E-01 |
|  | NTD | 1.7 | 0.05 | 1.7 | 0.36 | 4.E-01 |
| | ΔDBS | 1.1 | 0.09 | 2.3 | 0.68 | 1.E-03 |
| ***dnaA***<br>**mRNA** | FL | 0.51 | 0.02 | 0.71 | 0.45 | 2.E-01 |
|  | NTD | 1.5 | 0.04 | 2.0 | 0.35 | 6.E-03 |
| | ΔDBS | 0.86 | 0.04 | 2.5 | 0.49 | 4.E-09 |
| **9S**<br>**pre-rRNA** | FL | 2.2* | 0.08 | 2.5 | 0.29 | 6.E-03 |
|  | NTD | 2.5* | 0.25 | 2.2 | 0.29 | 1.E+00 |
| | ΔDBS | 4.8* | 0.35 | 4.3 | 0.66 | 2.E-01 |

\*Data from Al-Husini *et al.* Mol Cell 2018
<sup>#</sup>Stable 3' fragment removed from RNA-seq half-life calculation
<sup>&</sup> $\sigma$ calculated from standard deviation of linear regression
p-value calculated from a 1-tailed Z-test.

## Discussion

### BR-bodies exhibit selective permeability

Selective permeability is a hallmark for membrane bound organelles. Though bacteria generally lack membrane bound organelles, bacterial microcompartments (BMCs) such as carboxysomes have been shown to exhibit selective permeability to facilitate the carbon fixation pathway (Kerfeld et al., 2018), however, BMCs exhibit a rather narrow species distribution. Biomolecular condensates are widespread across eukaryotic cells (Banani et al., 2017), yet their identification across bacteria has only recently been explored (Abbondanzieri and Meyer, 2019; Al-Husini et al., 2018; Monterroso et al., 2019; Wang et al., 2019). Overall, BR-body condensates exhibit selective permeability by allowing in degradosome proteins, mRNAs, sRNAs, and asRNAs, and by rejecting ribosomes and the nucleoid. Such ability to organize enzymes (degradosomes) and their substrates (mRNAs) into biomolecular condensates may provide a more broadly utilized mechanism for bacterial subcellular organization. Indeed, rubisco itself was recently shown to form a biomolecular condensate with the cyanobacterial protein CcmM which facilitates the assembly of the carboxysome shell (Wang et al., 2019). While BR-bodies and CcmM-Rubisco condensates are currently the only bacterial biomolecular condensates whose components have been directly shown to form liquid-like droplets in physiological conditions (Al-Husini et al., 2018; Wang et al., 2019), the cell division protein FtsZ and its inhibitor SlmA were shown to form condensates upon addition of crowding reagents (Monterroso et al., 2019). In addition, other bacterial proteins such as the polar protein scaffold PopZ (Lasker et al., 2018; Zhao et al., 2018) and the nucleoid associated protein DPS (Janissen et al., 2018) have been shown to act as selectively permeable scaffolds, yet biomolecular condensation has not been directly observed.

Across domains of life, there is a well-known antagonistic relationship between the processes of mRNA translation and mRNA decay. Recent data have suggested that translation initiation and elongation can both impact mRNA decay (Chan et al., 2018; Presnyak et al., 2015), and translating ribosomes were found to be physically occluded from P-bodies and stress granules (Hubstenberger et al., 2017; Reineke et al., 2012). Here we showed that BR-bodies physically exclude ribosomes, providing a physical separation of mRNA decay and translation in bacteria (Al-Husini et al., 2018). The exclusion of ribosomes from BR-bodies may also explain why long poorly translated mRNAs are enriched BR-body substrates. While eukaryotic RNA silencing machinery is found in P-bodies and stress granules (Hubstenberger et al., 2017; Jain et al., 2016), sRNAs were found to be enriched in BR-bodies (Fig 3A) and are known to base pair with mRNAs often through the RNA chaperone Hfq (Santiago-Frangos and Woodson, 2018; Vogel and Luisi, 2011). Indeed, Hfq bound RNAs identified by HITS-CLIP (Assis et al., 2019) including both mRNAs and sRNAs are enriched in BR-bodies (Fig S6, Table S1), suggesting mRNA silencing may be influenced by BR-body localization. Cis-encoded asRNAs were also found to be enriched in BR-bodies (Fig 3) similar to stress granules where many short asRNAs associate through pairing to their complimentary mRNAs (Van Treeck et al., 2018). In bacteria asRNAs can be generated by internal promoters or by errors in rho-dependent transcription termination, and interestingly Rho protein was found to associate with RNase E when cells are grown at cold temperature (Aguirre et al., 2017), suggesting a link between these two processes. Importantly, mRNAs are the only class of substrate whose decay is globally stimulated by RNase E, while sRNAs and asRNAs have similar median half-lives in the ASM (Fig 1B, Table S1).

Most highly structured ncRNAs involved in translation-related processes (RNase P, tRNAs, SRP RNA, tmRNA, and rRNA) are highly depleted from BR-bodies. It is possible that the high level of secondary structure and/or strong association with specific RNA binding proteins limits these RNAs from entering BR-bodies. Interestingly, tmRNA, which functions to rescue non-stop mRNAs, was found to form diffraction limited foci in *C. crescentus* (Russell and Keiler, 2009), and we find tmRNA to be highly depleted from BR-bodies (Fig 3A), suggesting that ribosome rescue occurs in distinct cytoplasmic bodies.

### BR-bodies stimulate mRNA decay entry and prevent intermediate release

Despite broad identification of biomolecular condensates across organisms and cell types, the functional mechanisms of biomolecular condensates on their internal biochemical processes remains poorly understood. The high concentration of reactants and enzyme within the condensate has been proposed to have a stimulatory effect. This is true for the innate immune signaling protein cGAS, where condensation with DNA has a stimulatory effect on the rate of cGAMP production (Du and Chen, 2018). Similarly, condensate stimulation of actin polymerization has been shown for Arp2/3 N-WASP condensates (Banjade and Rosen, 2014; Li et al., 2012). In other cases, condensates have been proposed to allow substrate selectivity by partitioning only certain substrates together with the enzyme within the condensate (Banani et al., 2017).

In BR-bodies, the organization of the mRNA decay factors into a biomolecular condensate appears to perform multiple biochemical roles. BR-body condensation is stimulated upon the presence of untranslated mRNA (Fig 7) (Al-Husini et al., 2018). The NTD mutant of RNase E which cannot form condensates is slow in the initial mRNA cleavage step by RNase E (Fig 6) (Al-Husini et al., 2018), suggesting condensation stimulates initial cleavage. The stimulation of the initial cleavage rate is likely due to the condensate and not disruption of the RNA degradosome as a mutant lacking all degradosome binding sites (ΔDBS) in the IDR was still able to accelerate initial RNase E cleavage (Table 1) (Al-Husini et al., 2018). BR-body accelerated initial mRNA cleavage is likely conserved across bacteria as *E. coli* RNase E mutants that disrupt foci formation through IDR deletion or through deletion of a critical inner-membrane attachment helix required for foci formation both lead to slower global rates of mRNA decay (Hadjeras et al., 2019; Lopez et al., 1999; Strahl et al., 2015). Interestingly, the RNase E ΔDBS mutant lacking binding sites for exoribonucleases PNPase and RNase D was also found to have a lag between initial cleavage and decay of smaller RNA fragments suggesting a build-up of mRNA decay intermediates (Fig 6, Table 1). A similar buildup of mRNA decay intermediates is observed in *E. coli* when the conserved degradosome component PNPase is mutated (Coburn et al., 1999; Khemici and Carpousis, 2004; Morita et al., 2004). The accumulation of decay intermediates may explain why the ΔDBS strain grows more slowly (Fig S9) (Al-Husini et al., 2018). This suggests that another biochemical function of BR-bodies is to prevent the buildup of toxic mRNA decay intermediates (Fig 7). By spatially coordinating the assembly of poorly translated mRNAs and RNA decay enzymes into BR-bodies, bacteria can both accelerate the endo and exonuclease reaction rates and simultaneously prevent premature release of toxic reaction intermediates.

## Materials and Methods

### *Caulobacter crescentus* cell growth

All *Caulobacter crescentus* strains used in this study were derived from the wild-type strain NA1000 (Evinger and Agabian, 1977), and were grown at 28°C in peptone-yeast extract (PYE) medium or M2 minimal medium supplemented with 0.2% D-glucose (M2G) (Schrader and Shapiro, 2015). When appropriate, the indicated concentration of vanillate (500μM), xylose (0.2%), gentamycin (0.5 μg/mL), kanamycin (5 μg/mL), chloramphenicol (2 μg/mL), spectinomycin (25 μg/mL), and/or streptomycin (5 μg/mL) was added. Strains were analyzed at mid-exponential phase of growth (OD 0.3-0.6). Optical density was measured at 600 nm in a cuvette using a Nanodrop 2000C spectrophotometer. Replacements strains containing a xylose inducible copy of RNase E and a vanillate inducible RNase E variant were first grown in media containing xylose overnight, then washed 3 times with 1mL growth media, and resuspended in growth media containing vanillate, diluted, and grown overnight. Log-phase cultures were then used for the downstream experiment. **Ribosome occlusion:** JS348 and JS350 cells were grown overnight in PYE-Gent-Spec in 3 dilutions. Log-phase cells were split into two tubes and either treated with 10% Ethanol for 10 min. or left untreated as indicated. 1μL of the cells under each condition was spotted on a M2G 1.5% agarose pad and imaged using YFP, CFP, and TX-Red filter cubes as indicated. **Nucleoid occlusion:** JS299 cells were grown in PYE-Gent-Kan media containing 0.2% xylose overnight. The next day, the log-phase cells were washed 3 times with PYE media, and used to inoculate PYE-Gent-Kan media containing 0.5mM vanillate, diluted, and grown overnight. Log-phase cells were treated with 5 ng/µL DAPI for 15 minutes (cultures were covered with foil while shaking) and spotted on a M2G 1.5% agarose pad and imaged using both YFP and DAPI filter cubes.

### *E. coli* cell growth

*<u>E. coli</u>* strains were grown at 37°C and cultured in LB medium (L3522, Sigma), supplemented with the indicated concentration of kanamycin (30 μg/mL) or ampicillin (50 μg/mL). For induction, BL21 DE3 cells were induced with isopropyl-3-Dthiogalactopyranoside (IPTG) (1μM) for two hours and TOP10 cells were induced with 0.0004% <u>arabinose</u> for one hour. Strains were analyzed at mid-exponential phase of growth (OD_600_ 0.3-0.6). Optical density was measured at 600 nm in a cuvette using a NanoDrop 2000C spectrophotometer. **Nucleoid occlusion *(E. coli)*: JS230 and JS231 cells (TOP10)** were grown at 37°C in LB medium supplemented with 50 μg/mL ampicillin. For induction, log-phase cells were induced with 0.0004% arabinose 1 hour. After induction, cells were treated with 5ng/µl DAPI for 15 minutes before spotting on 1.5% agarose pad and imaging.

### *In vitro* droplet assembly assay

The RNase E CTD was purified as described in (Al-Husini et al., 2018). Cy5 dye-labeled RNAs were generated by *in vitro* transcription using a 9:1 ratio of UTP:aminoallyl-UTP (9S and tRNA samples (t Hoen et al., 2003)) followed by conjugation of a Cy5 NHS ester or by 3’ ligation of Cy5 pCp using T4 RNA ligase 1 (yeast total RNA, Poly-A (England et al., 1980)). Reactions were performed as described in (Al-Husini et al., 2018), where 12.4 µM RNase E CTD was incubated with the indicated RNA for 1-hour before imaging.

### Super-resolution imaging of BR-bodies and occlusion analysis

*C. crescentus* JS545 (L1-PAmCherry/RNaseE-eYFP) and JS546 (acnA-eYFP/RNase E-PAmCherry) cells were grown in liquid M2G media with 1.0 µg/mL gentamycin and 25 µg/mL spectinomycin to OD ∼0.6 and fixed at log phase using formaldehyde cross-linking with 1% fixation buffer (11% HCOH in 1x PBS at pH 7.3). The fixed cells were then spotted onto a pad of 1.5% agarose in M2G media and fiduciary 0.35 µm Fluoresbrite carboxylate YG beads were added at a concentration of 5.0 × 10^7^ spheres/mL. The cells were sandwiched between two coverslips and imaged on an Olympus IX71 inverted epifluorescence microscope with a 100× objective (NA 1.40). Each camera pixel corresponds to 49 nm × 49 nm in the sample.

Super-resolution microscopy was performed as described previously (Tuson and Biteen, 2015). Briefly, the eYFP fusions were imaged under 488-nm excitation (Coherent Sapphire 488-50) at a power density of ∼2.5 × 10^6^ µW/mm^2^. The PAmCherry fusions were photo-activated at 406 nm (Coherent Cube 406 laser) for 30 ms at 5.0 × 10^5^ µW/mm^2^. The PAmCherry was imaged under 561-nm excitation (Coherent Sapphire 561-50) at ∼2.5 × 10^6^ µW/mm^2^. The fluorescence emission was filtered with appropriate filters and imaged on a 512 × 512 pixel Photometrics Evolve electron-multiplying charge-coupled device (EMCCD) camera. Each channel was imaged sequentially. The PAmCherry was imaged first to avoid leakage of the eYFP signal (which extends to ∼600 nm) into the 561-nm channel.

### Super-Resolution Image Reconstruction

Single molecules were detected and fit as previously described using the SMALL-LABS algorithm (Isaacoff et al., 2019). In these fixed cells, individual molecules were fluorescent for multiple frames without moving. We therefore avoided over-counting single molecules with a spatio-temporal filter (Bayas et al., 2018): for each localized molecule, we fit the probability of finding an additional localization within 1.5 times the average 95% confidence interval of the fit over time to a biexponential decay, and used the short decay time as a temporal threshold. All single-molecule detections within 1.5 times the average 95% confidence interval and within the calculated temporal threshold were therefore combined as a single localization. We additionally avoided false positives by including only molecule detections that were tracked for at least five frames. Images were reconstructed as a histogram of number of fits in each 25 nm × 25 nm pixel on a grid, and then a Gaussian filter of 25 nm width was applied to the image to account for localization precision. To determine colocalization, RNase E foci were identified within these super-resolution histograms as collections of 25 nm × 25 nm pixels with counts greater than the average plus two standard deviations. A 1-pixel padding was added around each focus.

### Fluorescence In Situ Hybridization (FISH)

5 mL of cells were grown in M2G medium in a 28 °C shaker incubator in the presence of the appropriate antibiotics. For JS38 and JS299 the growth medium was supplied with 0.2% xylose, 0.5µg/mL gentamycin, and 5µg/mL kanamycin. Next day, the 5 mL overnight cultures were washed 3 times with M2G media and used to inoculate 25 mL M2G medium (in 3 dilutions) containing 500µM vanillate, 0.5µg/mL gentamycin, and 5µg/mL kanamycin and grown for 8 hours. JS403 and JP469 cells were grown in M2G medium with 0.5µg/mL gentamycin. For all the strains, log-phase cultures were then fixed with 7.5% para-formaldehyde (Sigma) for 15 min at 28 °C followed by incubation at 4 °C for 30 minutes. The fixed cells were harvested by centrifugation (11,000xg /3 min.). The cell pellets were washed 2 times in ice-cold 1% PBS (140mM NaCl, 3mM KCl, 8mM sodium phosphate, and 1.5mM potassium phosphate [PH 7.5]). The cell pellets were resuspended in ice-cold 100% ethanol and stored at -20°C for less than 1 week. When ready to proceed, the cells were pelleted at 4 °C (2655 × g for 5 minutes) and washed 3 times with GTE buffer (50 mM glucose, 20mM tris-HCl PH7, and 10mM EDTA). The cells were lysed with 4000U of RNase-free lysozyme (Lucigen) at 37 °C for 2 hours. Permeabilized cells were pelleted again and resuspended in 100 µL of hybridization buffer (Stellaris RNA FISH) containing 1µL of the *rsaA* FISH custom Assay DNA probes (Stellaris) or were RNase A treated prior to probing as a control. FISH probes with fluorescein Dye were used for probing *rsaA* of JS 403, and JP369 mCherry-popZ cells. FISH probes with Quasar 670 Dye were used for probing *rsaA* of JS38 and JS299 cells. The samples were incubated with the probes in the dark at 42 °C overnight. Next day, the cells were pelted and washed in wash buffer A (Stellaris RNA FISH) at 42 °C for 30 minutes. The cells were pelleted and washed with 1 mL of wash buffer B (Stellaris RNA FISH) for 5 minutes at 42 °C. the cells were pelleted and resuspended in GTE buffer supplied with 0.1% TX100. The suspended cells were spotted on a polylysine coated microscope slide, dried, washed 3 times with 1% PBS and air-dried again. A 5 µL drop of mounting medium (VECTASHIELD) was added, and the slides were covered with coverslips and the samples were imaged with a Nikon Eclipse NI-E with CoolSNAP MYO-CCD camera and 100x Oil CFI Plan Fluor (Nikon) objective, driven by Nikon elements software using appropriate filter cubes (Cy5, GFP, Texas Red). Using microbeJ (Ducret et al., 2016) the fluorescent foci were identified using the “maxima” function in microbeJ with “foci” selected as shape, with tolerance and Z-score parameters tuned for each image. Aberrant foci with area < 0.01 μm^2^ and length > 1 μM were removed, and the segmentation option was used to split adjoined foci.

### Visualization of the rsaA mRNA with the MS2 system

Strain JS287 containing a non-dimerizable mutant of the Ms2 coat protein fused to mCherry at the vanillate locus and an array of 96 Ms2 RNA hairpins (gift of Ido Golding) fused to the 3’ end of the *rsaA* gene were grown into log phase and induced with 0.5 mM vanillate for four hours and imaged on an M2G agarose pad. As a control, (JS25) lacking the integrated hairpins was imaged where no fluorescent foci were detected (Fig S10). Fluorescent foci were identified using microbeJ (Ducret et al., 2016) using the “maxima” function in microbeJ with “foci” selected as shape, with tolerance and Z-score parameters tuned for each image. Aberrant foci with area < 0.01 μm^2^ and length > 1 μM were removed, and the segmentation option was used to split adjoined foci. To measure colocalization of the *rsaA* mRNA with BR-bodies, we manually scored each mRNA focus for overlapping an RNase E focus or not. A minimum of 100 foci per strain were used for this analysis.

### Occlusion analysis of fluorescent microscopy images

Multi-channel images of test molecules with a BR-body protein fusion (either RNase E or aconitase) were aligned using the “align image by line ROI” plugin in FIJI. Next, fluorescent foci were identified using the “maxima” function in microbeJ with foci selected as shape, with tolerance and Z-score parameters tuned for each image. Aberrant foci with area < 0.01 μm^2^ and length > 1 μM were removed, and the segmentation option was used to split adjoined foci. For each focus, we manually drew a line-slice across the focus and recorded the fluorescence intensity of each channel, then ran a pearson correlation function on the intensities of each channel. In the case of the RNase E NTD expressed in *E. coli* where no foci form, the density of the nucleoid by DAPI intensity was used for the line slice instead. Resulting correlation coefficients were then reported for each focus, with a correlation coefficient of 1 representing perfect correlation, 0 representing no correlation, and -1 representing perfect anti-correlation (Fig S5). As a positive control we imaged (acnA-mCherry/RNE-eYFP) which both go into BR-bodies (Al-Husini et al., 2018), as an uncorrelated control we examined (AcnA-mCherry/RNEAconBS-YFP) where RNEAconBS-YFP still forms foci but AcnA-mCherry is diffuse across the cytoplasm (Al-Husini et al., 2018), and as a negative control we imaged (mCherry-PopZ/RNE-msfGFP) which the polar protein PopZ matrix is known to occlude ribosomes from the cell pole (Bowman et al., 2008) which also occludes the RNE-msfGFP protein. A minimum of 30 foci from 30 different cells were selected for each correlation distribution reported. Two-tailed T-tests with uneven variance were used to analyze statistical significance.

### Colony size analysis

Colonies of bacterial strains harboring the wild type (JS38), NTD (JS221), or ΔDBS (JS233) RNase E variants were grown on PYE gent-kan-vanillate plates and imaged on a gel imager with a black background. The colonies were thresholded manually in FIJI, and then the pixel density in the analyze-particle function were reported for a minimum of 5 colonies. For each strain, the average colony size was reported relative to the JS38 strain. Two-tailed T-tests with uneven variance were used to analyze statistical significance.

### BR-bodies Enrichment by Differential Centrifugation

5mL of JS299 cells were grown overnight in PYE medium supplied with 0.2% xylose, 0.5µg/mL gentamycin, and 5µg/mL kanamycin. The overnight cultures were then washed 3 times with PYE growth media and used to inoculate 30 mL of PYE medium supplied with 500µM vanillate, 0.5µg/mL gentamycin, and 5µg/mL kanamycin. The cells were grown overnight and then pelleted at 11,000xg for 5 min. The cell pellet was resuspended in 2.5 mL of lysis buffer (35mM NaCl, 20mM Tris-HCl-pH 7.4, 1mM β-mercaptoethanol, one tablet EDTA-free protease inhibitor (roche) per 25 mL of buffer, 1U/mL Superase IN, and 10U/ml RNase-free DNase I). The cell suspension was flash frozen dropwise in liquid nitrogen before lysis in a mixer-mill. After collecting a small scoop of the frozen lysate for the whole cell lysate sample, the cell lysate was spun at 2000xg for 5 min. to clear membranes. The pellet was resuspended in 200 µL of lysis buffer and the samples were spun again at 10,000xg for 10 min. The resulting pellet was resuspended again in 200 µL of lysis buffer and subjected to another spin at 20,000xg for 10 minutes. The BR-bodies enriched pellet was resuspended in 200 µL of lysis buffer and RNA was extracted from both whole cell lysate and BR-bodies enriched fractions. RNA extraction was performed by adding 1mL of 65°C Trizol to the samples and incubating at 65°C for 10 min, then 200μL of chloroform was added and incubated for 5 min at room temperature. The samples were then spun at max speed in a microcentrifuge for 10 min at room temperature and the aqueous layer was removed and incubated with 700μL of isopropanol. The samples were then precipitated at −20°C for 1 hour, spun at 20,000 g for 1 hour at 4°C, and washed three times with 80% ethanol. The pellet was air dried and resuspended in 10mM Tris pH 7.0. RNA-seq library construction was performed as described in (Aretakis et al., 2018) using 1.0 µg of total RNA. Raw sequencing data is available in the NCBI GEO database with accession number GSE133522.

### BR-body purification using HA-ASM pulldown

5 mL cultures of HA-ASM (JS302) and the untagged ASM (JS299) were inoculated and grown overnight in PYE-Gent-kan-xylose. The next day, these cultures were used to inoculate a 40 mL culture containing PYE/Gent/kan/xylose media and were diluted to be grown overnight. The log phase cultures were pelleted by centrifugation and washed 3x with 15 mL each of PYE. The washed cells were used to inoculate 50 mL of PYE-gent-kan-van and the cells were grown for 8 hours. The cells were harvested by centrifugation 11000xg for 10 min, resuspended in 0.5 mL of lysis buffer (35mM NaCl, 20 mM Tris-HCl (7.4), 1mM BME, 1U/ml Superase In, 10U/5mL RNase-free Dnase I). The pellets were flash frozen drop-wise in liquid nitrogen and stored at -80 C. The cells were lysed using mixer-mill and the cell lysates were thawed and transferred into Eppendorf tubes. The samples were spun at 2000xg for 5 min to remove membranes. 500 µL of the supernatant was used for HA-affinity purification. The purification was carried out as following:

100 µL of anti-HA-beads were washed three times (in 1.7 mL Eppendorf tubes) with 1mL of HA bead wash buffer each (20 mM Tris-HCl (7.4), 35mM NaCl, 1mM BME, 1U/ml Superase In). The cell lysate samples were added to the washed beads and incubated for 1 hour at 4 C on a nutator. The samples were spun at 12,000xg for 10 seconds and the flow through samples were collected. The beads were washed 3x with 1mL HA bead wash buffer for 20 min each. 100 µL of the HA-peptide solution was then added to the beads and were incubated at 30 C for 20 min and the first eluates were collected (E1). E2 samples were collected by adding another 100 µL of the HA peptide solution to the beads and incubating at 30 C for another 20 min. The third elution was done by adding 200 µL of the HA peptide to the samples and transferring them to Eppendorf tubes, incubating at 30 C for 5 min and collect the samples by spinning at 12,000xg for 10 seconds. The collected fractions were analyzed by imaging and western blot using anti-RNase E antibody (Gift from Luisi lab).

### Analysis of BR-body enrichment

mRNA length was estimated from transcriptional units mapped from previous RNA-seq datasets (Schrader et al., 2014; Zhou et al., 2015). To exclude complexities relating to multi-gene operons, simple mRNAs were used for the analysis. Simple mRNAs were defined as those containing a single TSS and a single CDS. mRNA levels, translation levels, and translation efficiency data from cells grown in M2G were from (Schrader et al., 2014). 5’ UTR length was calculated from (Schrader et al., 2014; Zhou et al., 2015). Shine-Dalgarno affinity was taken from (Schrader et al., 2014). TAI was calculated for *C. crescentus* by utilizing the online tool version of the TAI calculator (Sabi et al., 2017). *C. crescentus* was selected as the organism, and a FASTA file was uploaded with sequences of all of *C. crescentus* protein coding genes. stAIcalc (Sabi et al., 2017) calculated a TAI value for each gene as the geometrical mean of the TAI value of the codons making up that gene. nTE was calculated as described in as in (Pechmann and Frydman, 2013). RNA-seq data used in the calculation was from *C. crescentus* collected in M2G minimal media (Schrader et al., 2014). Hfq HITS-CLIP RNAs used were from (Assis et al., 2019).

### RNA decay rates by RNA-seq and qRT-PCR

JS38, JS221, JS233, and JS299 cells were grown in M2G-kan-gent media containing xylose overnight. The next day, the log-phase cultures were washed 3 times with M2G media and resuspended in 20 mL M2G-kan-gent media with 500 µM vanillate and grown for 8 hours. 1 mL of log-phase (OD600 0.3-0.6 *Caulobacter crescentus* cells untreated (0 min) or treated with (200 µg/mL) rifampicin for the indicated time points (1, 3, and 9 min). At the indicated time point the samples were added immediately to 2 mL of RNAprotect Bacterial Reagent (QIAGEN), immediately vortexed, and incubated at room temperature for 5 min. The cells were pelleted at (20,000xg) for 1 min and resuspended in 1mL of 65 °C hot TRizol Reagent (Ambion) and incubated 65 °C for 10 min in a heat block. 200µL of chloroform were added to the samples and the tubes were incubated at room temperature for 5 min before spinning at (20,000xg) for 10 min. RNA samples were chloroform extracted once and precipitated using isopropanol (1x volume isopropanol, 0.1X volume 5M NaOAc pH 5.2) overnight at −80 °C. The RNA samples were spun at 20,000 × g at 4 °C for 1 hour, pellets were washed with 80% ethanol for 10 min, air dried, and resuspended in 10 mM Tris-HCl (pH 7.0). The RNA-Seq libraries were made using 5µg of total RNA samples, rRNA was removed by ribozero gram negative kit, and library construction was performed according to protocol (Aretakis et al., 2018). Raw sequencing data is available in the NCBI GEO database with accession number GSE133532. The qRT-PCR, reactions were performed using 200 ng of total RNA samples according to the NEB Luna universal one-step-qRT-PCR kit on a stratagene MX3000P qRT-PCR machine. qRT-PCR were performed with primers for 5SrRNA, 9SrRNA, *rne, ctrA, gcrA*, and *dnaA* genes.

To measure RNA-decay rates we performed linear curve fitting of the ln (fraction RNA remaining) at each time point of RNA extraction. For RNA-seq, the fraction remaining was calculated as the RPKM of each time point divided by the RPKM measured in the untreated 0’ sample. For qRT-PCR, the Ct was converted into amount of RNA using a standard curve, and the amount of RNA at each time points was divided by the amount of RNA measured in the untreated 0’ sample. Slopes of the linear curve fits were then converted into mRNA decay rates using the following equation (mRNA half-life=-ln(2)/slope). To reduce biases in mRNA half-life induced by rifampicin lag (Chen et al., 2015), we excluded multi-gene operons from half-life measurement analysis and focused only on simple mRNAs. Simple mRNAs were defined as those containing a single TSS and a single ORF. For bulk mRNA half-life calculations of RNA-seq data the fraction of all mRNA reads was compared to the total fraction of reads which includes a majority of stable tRNA reads.

### Strain and Plasmid Construction

#### JS403: NA1000 rne::rne-mCherry Gent^R^

The RNase E insert was digested by NdeI/KpnI from pRNE-YFPC-1 and ligated to NdeI/KpnI digested pChyC-4. Resulting plasmids were transformed into *E. coli* and selected on LB gent plates, miniprepped, and sequence verified. Resulting pRNE-ChyC-4 was transformed into NA1000 cells, selected on PYE-gent plates, and verified by fluorescence microscopy.

#### JS25: NA1000 *vanA::Ms2(V75E/A81G)-mCherry* Spec^R^ Strp^R^

pVMs2(V75E/A81G)-mCherryC-1 plasmid was generated by IDT as a gblock fragment. The gblock fragment was assembled into pVChyC-1 (cut with NdeI and KpnI) via Gibson assembly. The insert was sequence verified then transformed into NA1000 cells via electroporation and selected on PYE spec/str. Resulting colonies were screened for vanillate inducible mCherry expression by fluorescence microscopy.

##### MS2 double mutant gblock

GCGAGGAAACGCATATGATGGCTTCTAACTTTACTCAGTTCGTTCTCGTCGACAATGGCGGAACTGGCGACGTGACTGTCGCCCCAAGCAACTTCGCTAACGGGGTCGCTGAATGGATCAGCTCTAACTCGCGTTCACAGGCTTACAAAGTAACCTGTAGCGTTCGTCAGAGCTCTGCGCAGAATCGCAAATACACCATCAAAGTCGAGGTGCCTAAAGTGGCAACCCAGACTGTTGGTGGAGAGGAGCTTCCTGTAGCCGGCTGGCGTTCGTACTTAAATATGGAACTAACCATTCCAATTTTCGCTACGAATTCCGACTGCGAGCTTATTGTTAAGGCAATGCAAGGTCTCCTAAAAGATGGAAACCCGATTCCCTCAGCAATCGCAGCAAACTCCGGCATCTACGGTACCTTAAGATCTCG

#### JS 430: NA1000 *vanA::Ms2(V75E/A81G)-mCherry rsaA::rsaA-96 array* Kan^R^ Spec^R^ Strp^R^

This strain was constructed at multiple steps. First, The Ms2 96 array was moved into pFlagC-2 by Gibson reaction. This was done by PCR amplifying the array from BAC2(P(lac,ara)- mRFP1 - 96bs) plasmid (Golding and Cox, 2004) with the following Gibson primers:

1. j5_00268_(Ms2array)_forward CTCCGGAGAATTCCGATTAGCTGCGCATCCCCC
2. j5_00269_(Ms2array)_reverse GGACAAAAACAGAGAAAGGAAACGACAGAGGCACCGGTCCGAC

The amplified 96 array fragment was gel purified and Gibson Ligated into pFlagC-2 plasmid (Thanbichler et al., 2007) amplified with the following Gibson primers:

1. j5_00270_(pflgc2EcoRIAgeI)_forward GGAAACGACAGAGGCACCGGTCCGACTACAAGGATGACG
2. j5_00271_(pflgc2EcoRIAgeI)_reverse CCTTAAGATCTCGAGCTCCGGAGAATTCCGATTAGCTGCGC

The Gibson ligation reaction was transformed into *E. coli* top10 cells and selected on LB-kan plates. Resulting kan^R^ colonies were then screened by PCR for the insert, and then verified by sanger sequencing(genewiz) pFlagC-2-96array. Second, he last 621 bp of *rsaA* gene was cloned into pFlagC-2-96array. This was done by PCR amplification of the *rsaA* fragment using the following primers:

1. rsaA F KpnI ATAAGGTACCCTGAACCTGACCAACACCGG
2. rsaA R EcoRI ATATTAGAATTCTTAGGCGAGCGTCAGGACTTCG

The PCR amplified fragment was gel purified, cut with Kpn1/EcoR1 and ligated to Kpn1/EcoR1 cut pFlagC-2-96array. The ligation was transformed into *E. coli* top10 cells and selected on LB-kan plates. Resulting kan^R^ colonies were then screened by PCR for the insert, and then verified by sanger sequencing(genewiz) making pFlagC-2-*rsaA*-96array. The purified plasmid was then transformed into NA1000 cells via electroporation and plated on PYE-kan plates.

Third, the pVMS2(V75E/A81G)-mCherryC-1 plasmid was transformed and cells were selected on PYE kan spec strep plates. Resulting colonies were screened for integration at the vanA locus (Thanbichler et al., 2007).

#### JS287: NA1000 *vanA::Ms2(V75E/A81G)-mCherry rsaA::rsaA-MS2 96 array rne*::rne-msfGFP Kan^R^ Spec^R^ Strp^R^ Gent^R^

JS430 cells were transduced with phage lysate from JS87 and selected on PYE gent plates. Resulting colonies were verified to have GFP expression.

#### JS348: NA1000 *L1::L1-eYFP acnA::acnA-mCherry* Gent^R^ Spec^R^

This strain was made by transducing JS290 cells(plasmid from Jared) with aconitase mcherry phage lysate from JS134 strain (Al-Husini et al., 2018) and the cells were plated on PYE-gent-spec-strp plates.

The growing colonies were restreaked three times, sequentially, on PYE-gent-spec-strp plates and validated by imaging

#### JS350: NA1000 *L1::L1-eYFP rne::rne-eCFP* Gent^R^ Spec^R^ Strp^R^

This strain was generated by Transducing JS255 cells with L1-eYFP phage lysate from JS 290 strain. The cells were plated on PYE-gent-spec-strp plates. The growing colonies were restreaked three times, sequentially, on PYE-gent-spec-strp plates and validated by imaging.

#### JS302: NA1000 *vanA::rne(HA-ASM_mut2_)-eYFP rne::pXrnessrAC* Gent^R^ Kan^R^

This strain was constructed by generating the pVRNE-HA-ASM_mut2_)-eYFPC-4 from pVRNE(ASM_mut_2)eYFPC-4 plasmid (Al-Husini et al., 2018) by site directed mutagenesis using the following primers:

1. HA add for TACCCGTACGACGTCCCGGACTACGCCCTGATCGACGCAGCACACG
2. HA add rev CATCTTCTTCGACATCATATGGTCGTC

First, the fragment was PCR amplified using the T4 PNK kinased oligos and using pVRNE(ASM mut2)YFPC-4 plasmid as a template. The resulting PCR reaction was gel purified and self-ligated. The ligation was transformed into *E. coli* TOP10 cells and selected on LB-gent plates. Resulting gent^R^ colonies were then verified by sanger sequencing (genewiz). Second the purified plasmid was transformed into NA1000 cells by electroporation and the transformants were selected by plating on PYE-Gent plates. Resulting colonies were grown in the presence and absence of vanillate and verified by fluorescence microscopy. Third, the NA1000 cells were transduced with phase lysate from JS8 cells (Al-Husini et al., 2018) and the cells were plated on PYE-Gent-Kan-xylose plates. The resultant colonies were patched on PYE-Gent-Kan plates with and without xylose to confirm the depletion phenotype.

#### JS545: NA1000 *L1::L1-PAmCherry rne::rne-eYFP* gent^R^ spec^R^ str^R^

Ribosomal protein L1 PAM fusion was generated by cloning the PAmCherry gene into pL1-YFPC-4 (Bayas et al., 2018). First NheI/AgeI digested pL1-YFPC-4 and PAmCherry insert was ligated to create pL1::PAMC-4. The resulting pL1::PAMC-4 plasmid was selected on LB-Kan plates and sequence verified. The plasmid was then transformed into NA1000 cells by electroporation, and the resulting gentR colonies were screened using fluorescence microscopy. Phage lysate from strain JS51 was then transduced into strain cells harboring pL1::PAMC-4 and selected on PYE-spec-gent plates. Resulting colonies were verified to express both fusions by fluorescence microscopy.

#### JS546: NA1000 *rne::rne-PAmCherry acnA::acnA-eYFP* gent^R^ spec^R^ str^R^

pRNE-YFPC-1 was first digested with NdeI/KpnI and ligated with pPAmCherryC-4 (NdeI/KpnI digested) and transformed into *E. coli* cells. Resulting gent^R^ colonies were screened for the insert and sequence verified yielding pRNE-PAmCherryC-4. pRNE-PAmCherryC-4 was transformed into NA1000 cells and selected on PYE gent plates. Gent resistant conlonies were verified to have RNE-PAmCherry expression by fluorescence imaging. Next cells were transduced with phage lysate from JS251 (Al-Husini et al., 2018) and selected on PYE gent-spec-str plates and expression of both fusions was verified by fluorescence microscopy.

## Supporting information

Table of oligonucleotides

Table of strains and plasmids

Table of key resources

## Author Contributions

JMS and WSC designed study. NA performed *in vivo* cell biology experiments, BR-body enrichment, and functional mRNA half-life experiments. NA and MAS performed MS2 mRNA visualization experiments. DTT performed *in vitro* droplet formation assays where RNAs were prepared by NSM. ZP, TZ, and JSB performed super-resolution microscopy and its analysis. JMS, MMB, and JRA assisted in global RNA half-life profiling analysis. JMS and OB performed diffraction limited occlusion image analysis. AG generated critical bacterial strains. All authors read the final manuscript.

## Acknowledgements

The authors thank Ben Luisi for α-RNase E antibody and Olke Uhlenbeck for critical feedback. The authors thank Wayne State University startup funds to JMS and University of Pittsburgh for startup funds to WSC. Research reported in this publication was supported by NIGMS of the National Institutes of Health under award numbers R35GM124733 to JMS and R21-GM128022 to JSB.

## Supplemental Figure Legends

**Figure S1. Global mRNA decay profiling of wild type and active site mutant variants of RNase E.** A.) Bulk mRNA decay rates for the wild type (JS38) and active site mutant (ASM) (JS299) RNase E variants. Bulk mRNA decay rate is calculated from the natural log of the fraction of RNA reads compared to all RNA reads (-Ln (2)/slope). B.) Log_2_ ratio of the mRNA half-life in the wild type divided by the mRNA half-life in the ASM.

**Figure S2. BR-body enrichment procedure**. A.) Flow chart of the BR-body enrichment procedure with differential centrifugation adapted from (Khong et al., 2018; Khong et al., 2017; Wheeler et al., 2017). Microscopy images of ASM-YFP were performed with pellets resuspended in the same volume as supernatant for comparison. White scale bar represents 25 µm. B.) HA-tagged RNase E ASM bodies do not remain intact during Immunoprecipitation (IP). Western blot of whole cell lysate (WCL), flow through (FT), and sequential Elutions (E1-E3) using anti-RNase E antibodies. Below, images of the pooled elution samples of mock IP of the ASM (JS299) from the IP procedure (left), HA-tagged RNase E ASM (JS302) from the IP procedure (∼3.5 hours total procedure time) (center) or the rapid differential centrifugation procedure (∼35 minute total procedure time) (right). No large bodies remained after HA-IP, while they were readily detected in the rapid differential centrifugation protocol. C.) Bioanalyzer traces of total RNA extracted from the WCL fraction (top) or from the enriched samples generated from differential centrifugation (bottom). Samples were collected from the ASM (JS299) strain.

**Figure S3. Determinants of BR-body enrichment for mRNAs**. A.-B.) Bar graphs of mRNA features separated by bins of BR-body enrichment. To reduce complexity arising from multi-gene operons, analysis was performed only on simple mRNAs defined as having only a single TSS and a single CDS. C.) Correlation coefficient of mRNA features related to BR-body enrichment.

**Figure S4. Determinants of BR-body enrichment for sRNAs**. A.) comparison of mRNA and sRNA enrichment by length. sRNAs are colored red, mRNAs are colored black. B.) Bar graphs of sRNA features separated by bins of BR-body enrichment.

**Figure S5. BR-body occlusion analysis**. A.) Line slice analysis of BR-body colocalization patterns. A line slice across the center of a foci in one channel were correlated with the intensity of the fluorescence in a separate channel with the possible scenarios shown on the left. On the right, control data for known proteins that colocalize with BR-bodies (AcnA-mCherry/RNE-YFP), contain no significant colocalization (AcnA-mCherry/RNE-AconBS-YFP), and a protein that occludes BR-bodies (PopZ-mCherry/RNE-msfGFP). A minimum of 30 foci were included in the analysis for each comparison. B.) correlation coefficients for translation related molecules. C.) Correlation coefficients for RNase E variants and the nucleoid.

**Figure S6. BR-body enrichment of Hfq associated RNAs**. Box plot of BR-body enrichment Hfq bound RNAs determined by HITS-CLIP (Assis et al., 2019).

**Fig. S7. Super-resolution co-localization of RNase E and aconitase and nucleoid occlusion of the NTD and CTD**. A.) Super-resolution images of aconitase-eYFP (magenta) and RNase E-PAmCherry (green). Cell outlines from phase image. Scale bar is 500 nm. Below, multiple cells of aconitase-eYFP/RNE-PAmCherry (left) and RNase E-eYFP/L1-PAmCherry (right). Each Image is 2 × 3.5 µm B.) *E. coli* strains harboring pBad copies of the CTD-eYFP or NTD-eYFP variants were induced with 0.0004% arabinose for 1 hour and imaged with DAPI and YFP filter cubes. Red line represents the location where the intensity of each channel displayed on the right is generated from.

**Fig. S8. Distribution of mRNA half-lives from the global profiling experiment**. Top, Histogram of the mRNA half-lives across transcripts in the the JS38 (blue), JS221 (orange), JS233 (yellow), and JS299 (green) strains. Bottom, histogram of the Log2 fold change (mut/wt) in mRNA half-lives for each mutant.

**Figure S9. Colony sizes of RNase E variant strains**. Relative colony sizes of RNase E replacement strains expressing wild type RNase E (JS38), NTD only (JS221), or the ΔDBS mutant (JS233). Cells were grown in PYE-vanillate-gentamycin and imaged in a gel imager with white light. Colony size was measured in imageJ by thresholding the image and using the analyze particle function.

**Figure S10. mRNA localization control experiments**. Ms2 labeling system controls. A.) Fraction of cells containing mRNA spots. Number of cells=186 *rsaA*, 213 no array, 418 rifampicin treated. B.) Time course of Ms2 foci upon rifampicin addition. Half-life of Ms2 foci calculated from ln(2)/slope. Number of cells analyzed at each timepoint= 2762 0min, 496 5min, 1468 9min, 1949 14min, 743 23min, 1044 60min. C.) mRNA FISH with NA1000 cells using the fluorescein FISH probes (left). mRNA FISH controls (right) using either RNase A pre-treatment of cells prior to FISH probing (top) or omitting probes (bottom) are shown.

## References

Abbondanzieri, E.A., and Meyer, A.S. (2019). More than just a phase: the search for membraneless organelles in the bacterial cytoplasm. Curr Genet.

Aguirre, A.A., Vicente, A.M., Hardwick, S.W., Alvelos, D.M., Mazzon, R.R., Luisi, B.F., and Marques, M.V. (2017). Association of the Cold Shock DEAD-Box RNA Helicase RhlE to the RNA Degradosome in Caulobacter crescentus. J Bacteriol 199.

Ait-Bara, S., and Carpousis, A.J. (2015). RNA degradosomes in bacteria and chloroplasts: classification, distribution and evolution of RNase E homologs. Mol Microbiol 97, 1021–1135.

Al-Husini, N., Tomares, D.T., Bitar, O., Childers, W.S., and Schrader, J.M. (2018). alpha-Proteobacterial RNA Degradosomes Assemble Liquid-Liquid Phase-Separated RNP Bodies. Mol Cell 71, 1027–1039 e1014.

Aretakis, J.R., Al-Husini, N., and Schrader, J.M. (2018). Methodology for Ribosome Profiling of Key Stages of the Caulobacter crescentus Cell Cycle. Methods Enzymol 612, 443–465.

Assis, N.G., Ribeiro, R.A., da Silva, L.G., Vicente, A.M., Hug, I., and Marques, M.V. (2019). Identification of Hfq-binding RNAs in Caulobacter crescentus. RNA Biol, 1–8.

Banani, S.F., Lee, H.O., Hyman, A.A., and Rosen, M.K. (2017). Biomolecular condensates: organizers of cellular biochemistry. Nat Rev Mol Cell Biol 18, 285–298.

Bandyra, K.J., and Luisi, B.F. (2018). RNase E and the High-Fidelity Orchestration of RNA Metabolism. Microbiology spectrum 6.

Banjade, S., and Rosen, M.K. (2014). Phase transitions of multivalent proteins can promote clustering of membrane receptors. eLife 3.

Bayas, C.A., Wang, J., Lee, M.K., Schrader, J.M., Shapiro, L., and Moerner, W.E. (2018). Spatial organization and dynamics of RNase E and ribosomes in *Caulobacter crescentus*. Proceedings of the National Academy of Sciences, In Press.

Bowman, G.R., Comolli, L.R., Zhu, J., Eckart, M., Koenig, M., Downing, K.H., Moerner, W.E., Earnest, T., and Shapiro, L. (2008). A polymeric protein anchors the chromosomal origin/ParB complex at a bacterial cell pole. Cell 134, 945–955.

Callaghan, A.J., Marcaida, M.J., Stead, J.A., McDowall, K.J., Scott, W.G., and Luisi, B.F. (2005). Structure of Escherichia coli RNase E catalytic domain and implications for RNA turnover. Nature 437, 1187–1191.

Chan, L.Y., Mugler, C.F., Heinrich, S., Vallotton, P., and Weis, K. (2018). Non-invasive measurement of mRNA decay reveals translation initiation as the major determinant of mRNA stability. eLife 7.

Chen, H., Shiroguchi, K., Ge, H., and Xie, X.S. (2015). Genome-wide study of mRNA degradation and transcript elongation in *Escherichia coli*. Mol Syst Biol 11, 781.

Clarke, J.E., Kime, L., Romero, A.D., and McDowall, K.J. (2014). Direct entry by RNase E is a major pathway for the degradation and processing of RNA in Escherichia coli. Nucleic Acids Res 42, 11733–11751.

Coburn, G.A., Miao, X., Briant, D.J., and Mackie, G.A. (1999). Reconstitution of a minimal RNA degradosome demonstrates functional coordination between a 3’ exonuclease and a DEAD-box RNA helicase. Genes Dev 13, 2594–2603.

Courchaine, E.M., Lu, A., and Neugebauer, K.M. (2016). Droplet organelles? EMBO J 35, 1603–1612.

Du, M., and Chen, Z.J. (2018). DNA-induced liquid phase condensation of cGAS activates innate immune signaling. Science 361, 704–709.

Ducret, A., Quardokus, E.M., and Brun, Y.V. (2016). MicrobeJ, a tool for high throughput bacterial cell detection and quantitative analysis. Nat Microbiol 1, 16077.

Ebersbach, G., Briegel, A., Jensen, G.J., and Jacobs-Wagner, C. (2008). A self-associating protein critical for chromosome attachment, division, and polar organization in *Caulobacter*. Cell 134, 956–968.

England, T.E., Bruce, A.G., and Uhlenbeck, O.C. (1980). Specific labeling of 3’ termini of RNA with T4 RNA ligase. Methods Enzymol 65, 65–74.

Eulalio, A., Behm-Ansmant, I., Schweizer, D., and Izaurralde, E. (2007). P-Body Formation Is a Consequence, Not the Cause, of RNA-Mediated Gene Silencing. Mol Cell Biol 27, 3970–3981.

Eulalio, A., Huntzinger, E., and Izaurralde, E. (2008). GW182 interaction with Argonaute is essential for miRNA-mediated translational repression and mRNA decay. Nat Struct Mol Biol 15, 346–353.

Evinger, M., and Agabian, N. (1977). Envelope-associated nucleoid from *Caulobacter crescentus* stalked and swarmer cells. J Bacteriol 132, 294–301.

Frohlich, K.S., Forstner, K.U., and Gitai, Z. (2018). Post-transcriptional gene regulation by an Hfq-independent small RNA in *Caulobacter crescentus*. Nucleic Acids Res 46, 10969–10982.

Golding, I., and Cox, E.C. (2004). RNA dynamics in live Escherichia coli cells. Proc Natl Acad Sci U S A 101, 11310–11315.

Hadjeras, L., Poljak, L., Bouvier, M., Morin-Ogier, Q., Canal, I., Cocaign-Bousquet, M., Girbal, L., and Carpousis, A.J. (2019). Detachment of the RNA degradosome from the inner membrane of Escherichia coli results in a global slowdown of mRNA degradation, proteolysis of RNase E and increased turnover of ribosome-free transcripts. Mol Microbiol 0.

Hammarlof, D.L., Bergman, J.M., Garmendia, E., and Hughes, D. (2015). Turnover of mRNAs is one of the essential functions of RNase E. Mol Microbiol 98, 34–45.

Hardwick, S.W., Chan, V.S., Broadhurst, R.W., and Luisi, B.F. (2011). An RNA degradosome assembly in Caulobacter crescentus. Nucleic Acids Res 39, 1449–1459.

Hubstenberger, A., Courel, M., Benard, M., Souquere, S., Ernoult-Lange, M., Chouaib, R., Yi, Z., Morlot, J.B., Munier, A., Fradet, M., et al. (2017). P-Body Purification Reveals the Condensation of Repressed mRNA Regulons. Mol Cell 68, 144–157 e145.

Hui, M.P., Foley, P.L., and Belasco, J.G. (2014). Messenger RNA degradation in bacterial cells. Annu Rev Genet 48, 537–559.

Isaacoff, B.P., Li, Y., Lee, S.A., and Biteen, J.S. (2019). SMALL-LABS: Measuring Single-Molecule Intensity and Position in Obscuring Backgrounds. Biophys J 116, 975–982.

Ivanov, P., Kedersha, N., and Anderson, P. (2019). Stress Granules and Processing Bodies in Translational Control. Cold Spring Harb Perspect Biol 11.

Jain, S., Wheeler, Joshua R., Walters, Robert W., Agrawal, A., Barsic, A., and Parker, R. (2016). ATPase-Modulated Stress Granules Contain a Diverse Proteome and Substructure. Cell 164, 487–498.

Janissen, R., Arens, M.M.A., Vtyurina, N.N., Rivai, Z., Sunday, N.D., Eslami-Mossallam, B., Gritsenko, A.A., Laan, L., de Ridder, D., Artsimovitch, I., et al. (2018). Global DNA Compaction in Stationary-Phase Bacteria Does Not Affect Transcription. Cell 174, 1188–1199 e1114.

Kerfeld, C.A., Aussignargues, C., Zarzycki, J., Cai, F., and Sutter, M. (2018). Bacterial microcompartments. Nat Rev Microbiol 16, 277.

Khemici, V., and Carpousis, A.J. (2004). The RNA degradosome and poly(A) polymerase of Escherichia coli are required in vivo for the degradation of small mRNA decay intermediates containing REP-stabilizers. Mol Microbiol 51, 777–790.

Khong, A., Jain, S., Matheny, T., Wheeler, J.R., and Parker, R. (2018). Isolation of mammalian stress granule cores for RNA-Seq analysis. Methods 137, 49–54.

Khong, A., Matheny, T., Jain, S., Mitchell, S.F., Wheeler, J.R., and Parker, R. (2017). The Stress Granule Transcriptome Reveals Principles of mRNA Accumulation in Stress Granules. Mol Cell 68, 808–820 e805.

Landt, S.G., Lesley, J.A., Britos, L., and Shapiro, L. (2010). CrfA, a small noncoding RNA regulator of adaptation to carbon starvation in *Caulobacter crescentus*. J Bacteriol 192, 4763–4775.

Lasker, K., von Diezmann, A., Ahrens, D.G., Mann, T.H., Moerner, W.E., and Shapiro, L. (2018). Phospho-signal flow from a pole-localized microdomain spatially patterns transcription factor activity. bioRxiv, 220293.

Lau, J.H., Nomellini, J.F., and Smit, J. (2010). Analysis of high-level S-layer protein secretion in *Caulobacter crescentus*. Can J Microbiol 56, 501–514.

LeCuyer, K.A., Behlen, L.S., and Uhlenbeck, O.C. (1995). Mutants of the bacteriophage MS2 coat protein that alter its cooperative binding to RNA. Biochemistry 34, 10600–10606.

Li, P., Banjade, S., Cheng, H.C., Kim, S., Chen, B., Guo, L., Llaguno, M., Hollingsworth, J.V., King, D.S., Banani, S.F., et al. (2012). Phase transitions in the assembly of multivalent signalling proteins. Nature 483, 336–340.

Lopez, P.J., Marchand, I., Joyce, S.A., and Dreyfus, M. (1999). The C-terminal half of RNase E, which organizes the Escherichia coli degradosome, participates in mRNA degradation but not rRNA processing in vivo. Mol Microbiol 33, 188–199.

Mohanty, B.K., and Kushner, S.R. (2016). Regulation of mRNA Decay in Bacteria. Annu Rev Microbiol 70, 25–44.

Montero Llopis, P., Jackson, A.F., Sliusarenko, O., Surovtsev, I., Heinritz, J., Emonet, T., and Jacobs-Wagner, C. (2010). Spatial organization of the flow of genetic information in bacteria. Nature 466, 77–81.

Monterroso, B., Zorrilla, S., Sobrinos-Sanguino, M., Robles-Ramos, M.A., Lopez-Alvarez, M., Margolin, W., Keating, C.D., and Rivas, G. (2019). Bacterial FtsZ protein forms phase-separated condensates with its nucleoid-associated inhibitor SlmA. EMBO Rep 20.

Morita, T., Kawamoto, H., Mizota, T., Inada, T., and Aiba, H. (2004). Enolase in the RNA degradosome plays a crucial role in the rapid decay of glucose transporter mRNA in the response to phosphosugar stress in Escherichia coli. Mol Microbiol 54, 1063–1075.

Ono, M., and Kuwano, M. (1979). A conditional lethal mutation in an Escherichia coli strain with a longer chemical lifetime of messenger RNA. J Mol Biol 129, 343–357.

Pechmann, S., and Frydman, J. (2013). Evolutionary conservation of codon optimality reveals hidden signatures of cotranslational folding. Nat Struct Mol Biol 20, 237–243.

Pitchiaya, S., Mourao, M.D.A., Jalihal, A.P., Xiao, L., Jiang, X., Chinnaiyan, A.M., Schnell, S., and Walter, N.G. (2019). Dynamic Recruitment of Single RNAs to Processing Bodies Depends on RNA Functionality. Mol Cell 74, 521–533.e526.

Presnyak, V., Alhusaini, N., Chen, Y.H., Martin, S., Morris, N., Kline, N., Olson, S., Weinberg, D., Baker, K.E., Graveley, B.R., et al. (2015). Codon optimality is a major determinant of mRNA stability. Cell 160, 1111–1124.

Protter, D.S., and Parker, R. (2016). Principles and Properties of Stress Granules. Trends Cell Biol 26, 668–679.

Reineke, L.C., Dougherty, J.D., Pierre, P., and Lloyd, R.E. (2012). Large G3BP-induced granules trigger eIF2alpha phosphorylation. Mol Biol Cell 23, 3499–3510.

Russell, J.H., and Keiler, K.C. (2009). Subcellular localization of a bacterial regulatory RNA. Proc Natl Acad Sci U S A 106, 16405–16409.

Sabi, R., Volvovitch Daniel, R., and Tuller, T. (2017). stAIcalc: tRNA adaptation index calculator based on species-specific weights. Bioinformatics 33, 589–591.

Santiago-Frangos, A., and Woodson, S.A. (2018). Hfq chaperone brings speed dating to bacterial sRNA. Wiley interdisciplinary reviews RNA 9, e1475.

Schrader, J.M., and Shapiro, L. (2015). Synchronization of Caulobacter crescentus for investigation of the bacterial cell cycle. J Vis Exp, e52633.

Schrader, J.M., Zhou, B., Li, G.W., Lasker, K., Childers, W.S., Williams, B., Long, T., Crosson, S., McAdams, H.H., Weissman, J.S., et al. (2014). The coding and noncoding architecture of the *Caulobacter crescentus* genome. PLoS Genet 10, e1004463.

Sheth, U., and Parker, R. (2003). Decapping and decay of messenger RNA occur in cytoplasmic processing bodies. Science 300, 805–808.

Shin, Y., and Brangwynne, C.P. (2017). Liquid phase condensation in cell physiology and disease. Science 357.

Strahl, H., Turlan, C., Khalid, S., Bond, P.J., Kebalo, J.M., Peyron, P., Poljak, L., Bouvier, M., Hamoen, L., Luisi, B.F., et al. (2015). Membrane recognition and dynamics of the RNA degradosome. PLoS Genet 11, e1004961.

Surovtsev, I.V., and Jacobs-Wagner, C. (2018). Subcellular Organization: A Critical Feature of Bacterial Cell Replication. Cell 172, 1271–1293.

t Hoen, P.A., de Kort, F., van Ommen, G.J., and den Dunnen, J.T. (2003). Fluorescent labelling of cRNA for microarray applications. Nucleic Acids Res 31, e20.

Thanbichler, M., Iniesta, A.A., and Shapiro, L. (2007). A comprehensive set of plasmids for vanillate- and xylose-inducible gene expression in Caulobacter crescentus. Nucleic Acids Res 35, e137.

Tien, M., Fiebig, A., and Crosson, S. (2018). Gene network analysis identifies a central post-transcriptional regulator of cellular stress survival. eLife 7.

Tuson, H.H., and Biteen, J.S. (2015). Unveiling the inner workings of live bacteria using super-resolution microscopy. Anal Chem 87, 42–63.

Van Treeck, B., Protter, D.S.W., Matheny, T., Khong, A., Link, C.D., and Parker, R. (2018). RNA self-assembly contributes to stress granule formation and defining the stress granule transcriptome. Proc Natl Acad Sci U S A 115, 2734–2739.

Vogel, J., and Luisi, B.F. (2011). Hfq and its constellation of RNA. Nat Rev Microbiol 9, 578–589.

Voss, J.E., Luisi, B.F., and Hardwick, S.W. (2014). Molecular recognition of RhlB and RNase D in the Caulobacter crescentus RNA degradosome. Nucleic Acids Res 42, 13294–13305.

Wang, C., Schmich, F., Srivatsa, S., Weidner, J., Beerenwinkel, N., and Spang, A. (2018). Context-dependent deposition and regulation of mRNAs in P-bodies. eLife 7.

Wang, H., Yan, X., Aigner, H., Bracher, A., Nguyen, N.D., Hee, W.Y., Long, B.M., Price, G.D., Hartl, F.U., and Hayer-Hartl, M. (2019). Rubisco condensate formation by CcmM in β-carboxysome biogenesis. Nature 566, 131–135.

Weber, S.C. (2017). Sequence-encoded material properties dictate the structure and function of nuclear bodies. Curr Opin Cell Biol 46, 62–71.

Wheeler, J.R., Jain, S., Khong, A., and Parker, R. (2017). Isolation of yeast and mammalian stress granule cores. Methods 126, 12–17.

Wheeler, J.R., Matheny, T., Jain, S., Abrisch, R., and Parker, R. (2016). Distinct stages in stress granule assembly and disassembly. eLife 5.

Zhao, W., Duvall, S.W., Kowallis, K.A., Tomares, D.T., Petitjean, H.N., and Childers, W.S. (2018). A circuit of protein-protein regulatory interactions enables polarity establishment in a bacterium. bioRxiv, 503250.

Zhou, B., Schrader, J.M., Kalogeraki, V.S., Abeliuk, E., Dinh, C.B., Pham, J.Q., Cui, Z.Z., Dill, D.L., McAdams, H.H., and Shapiro, L. (2015). The Global Regulatory Architecture of Transcription during the *Caulobacter* Cell Cycle. PLoS Genet 11, e1004831.

