## Supplementary figures and images for "BR-bodies provide selectively permeable condensates that stimulate mRNA decay and prevent release of decay intermediates"

### Fig S1

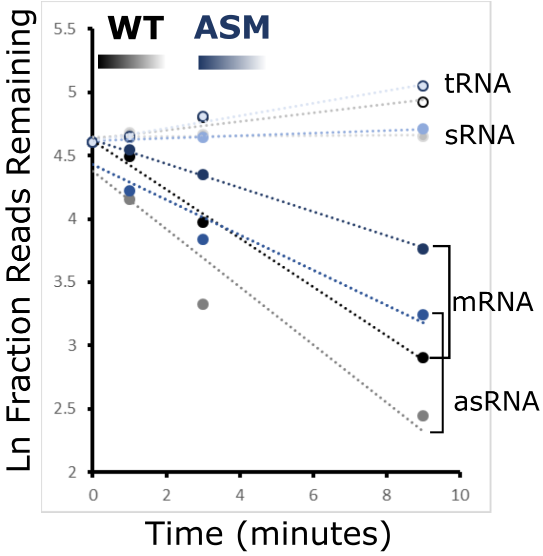

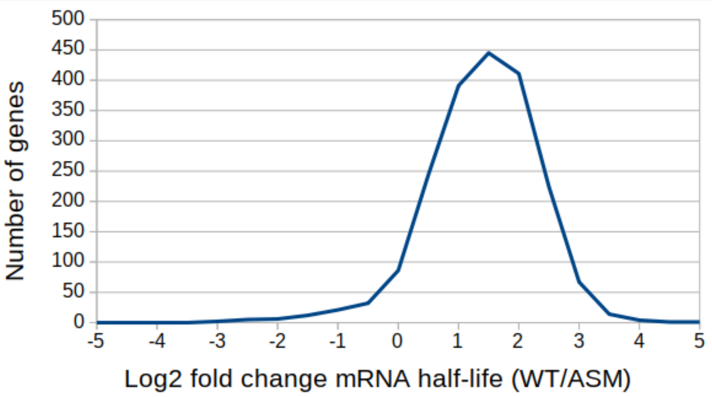

### Fig S2

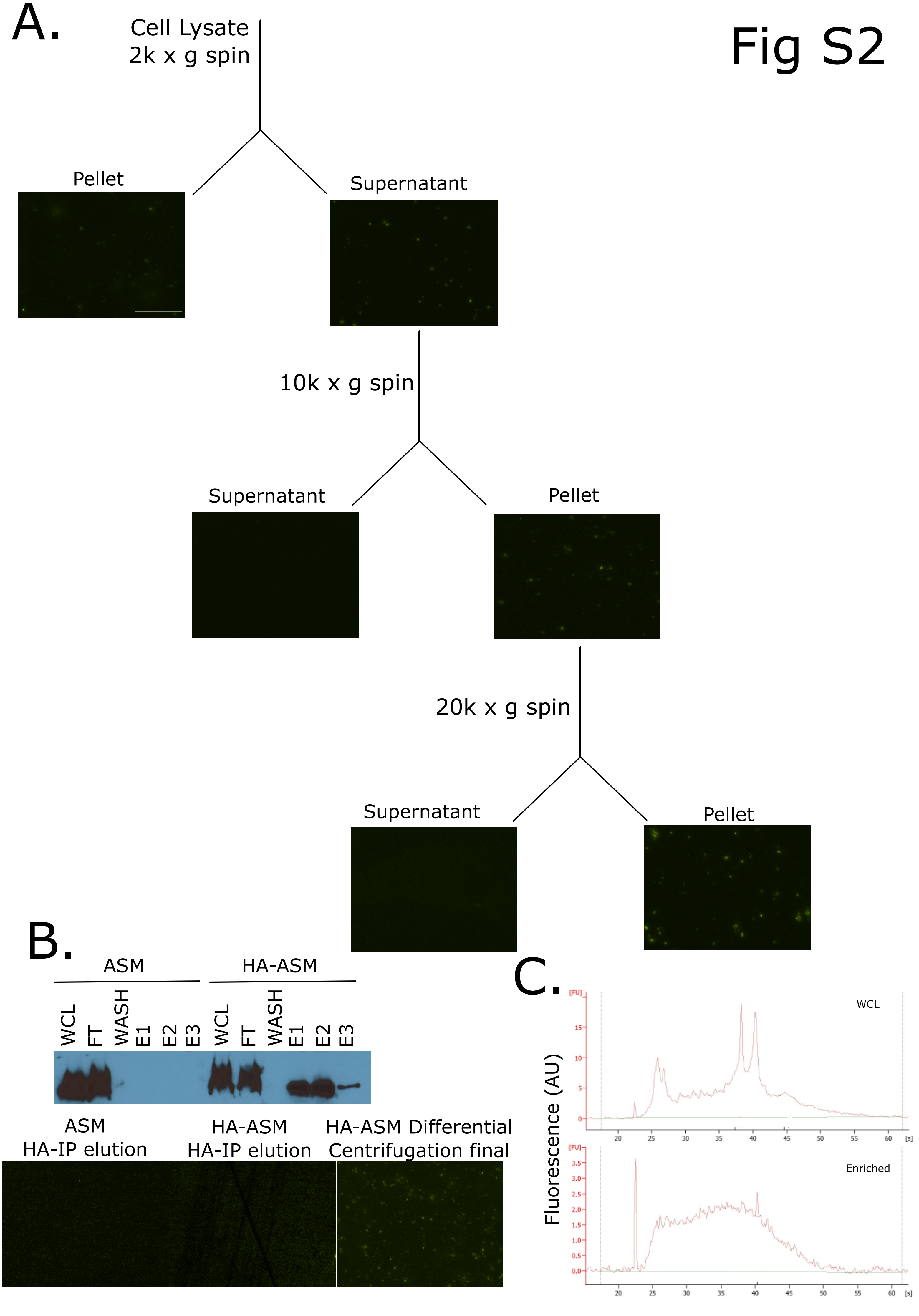

### Fig S4

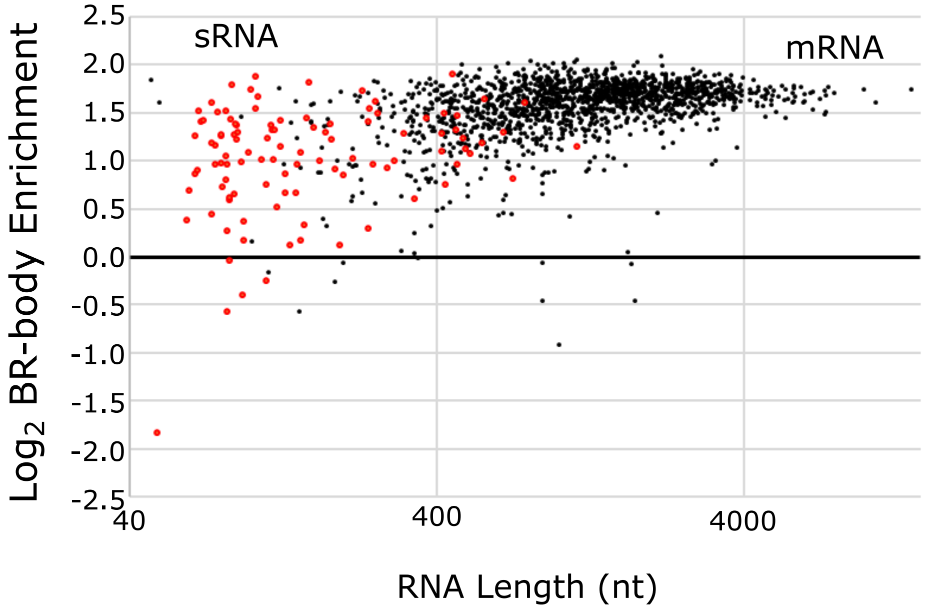

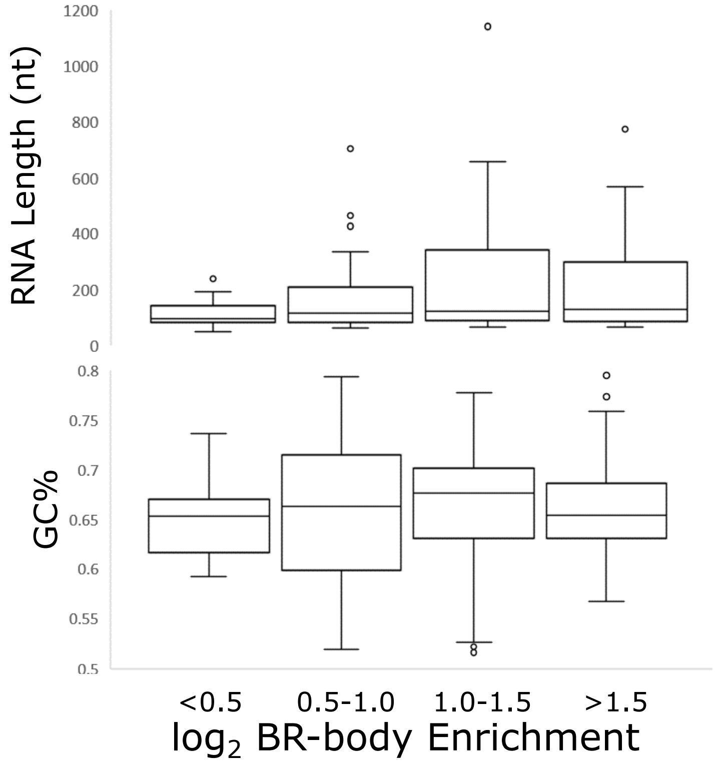

### Fig S5

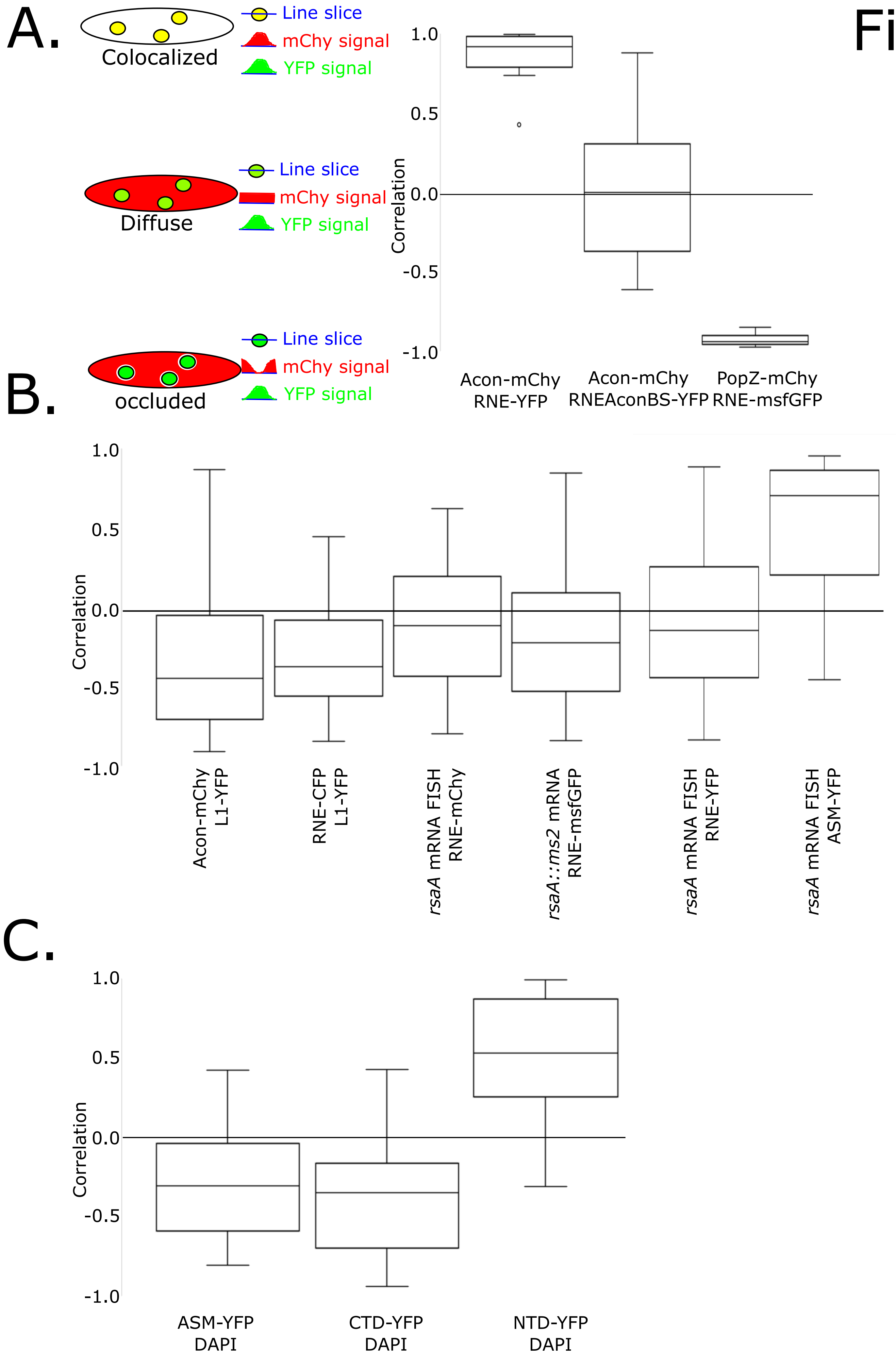

### Fig S6

Fig S6

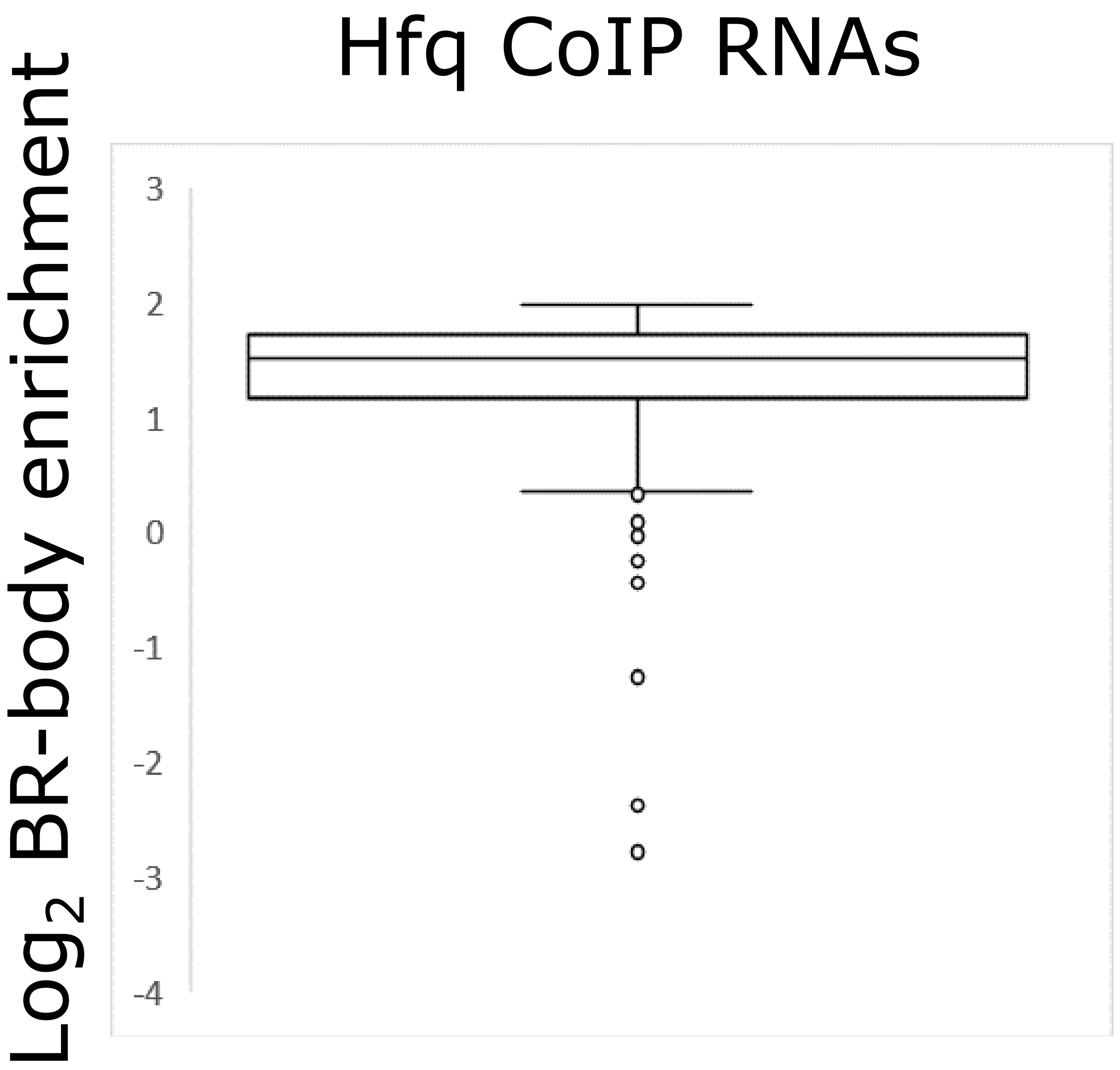

### Fig S7

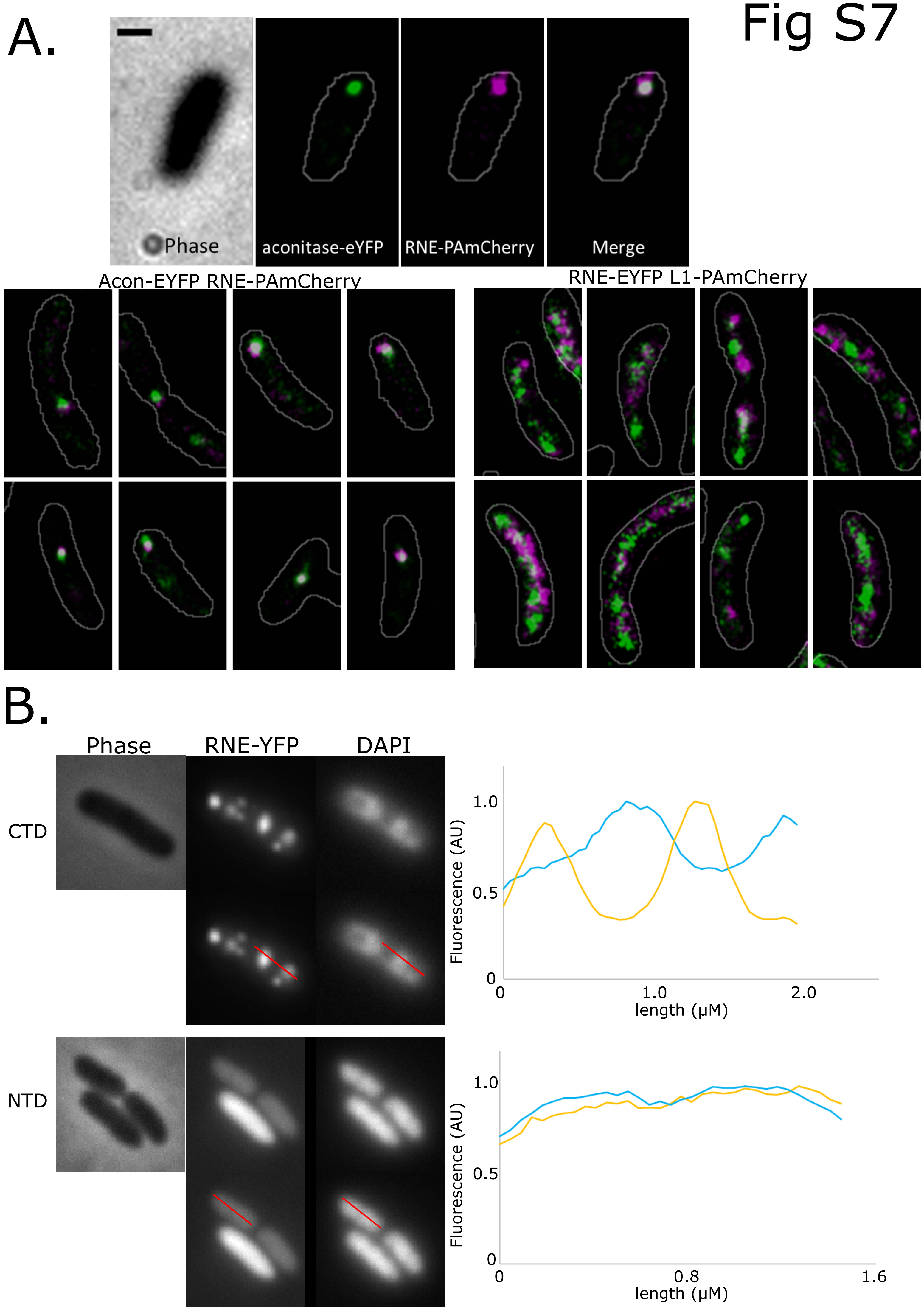

### Fig S8

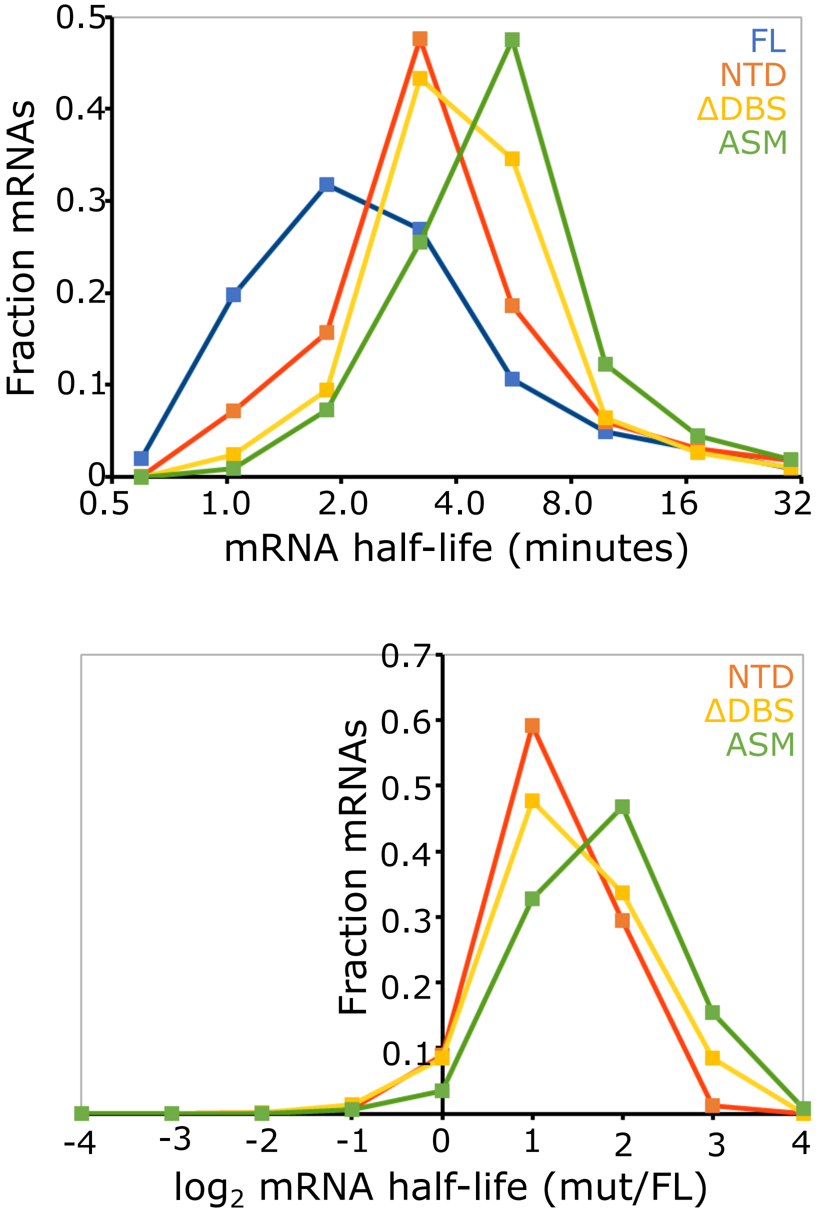

### Fig S9

Fig S9

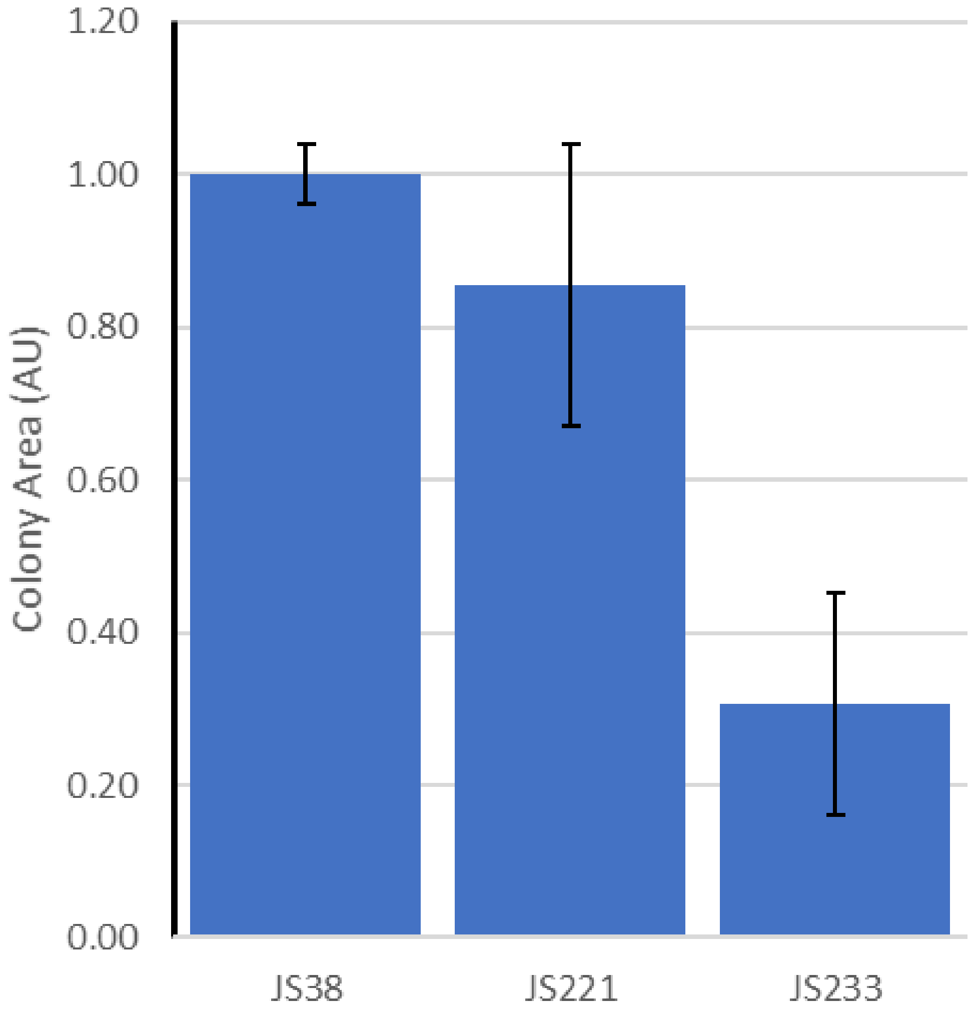

### Fig S10

#### Fig S10

##### MS2 rsaA mRNA labeling controls

A.

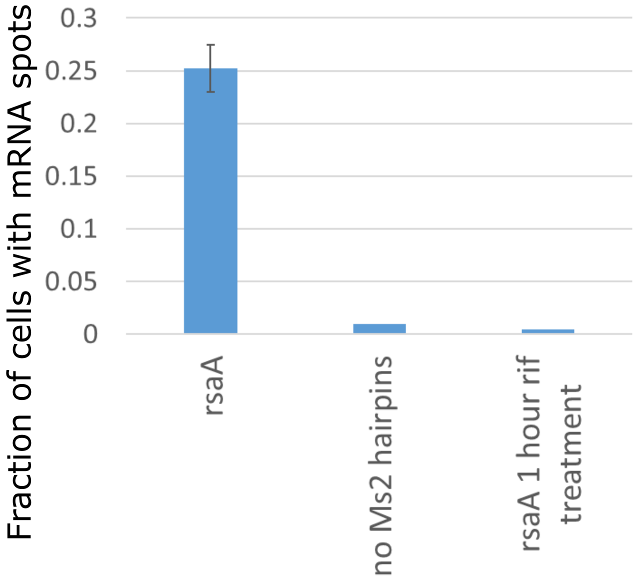


B.

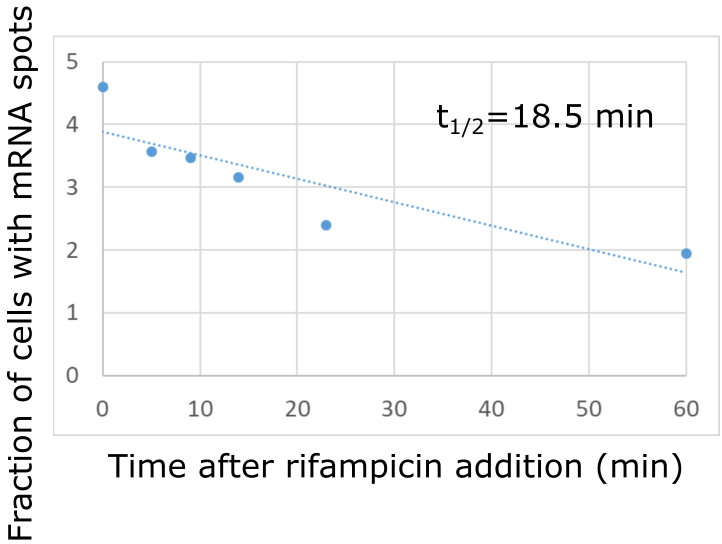


##### rsaA mRNA FISH controls

C.

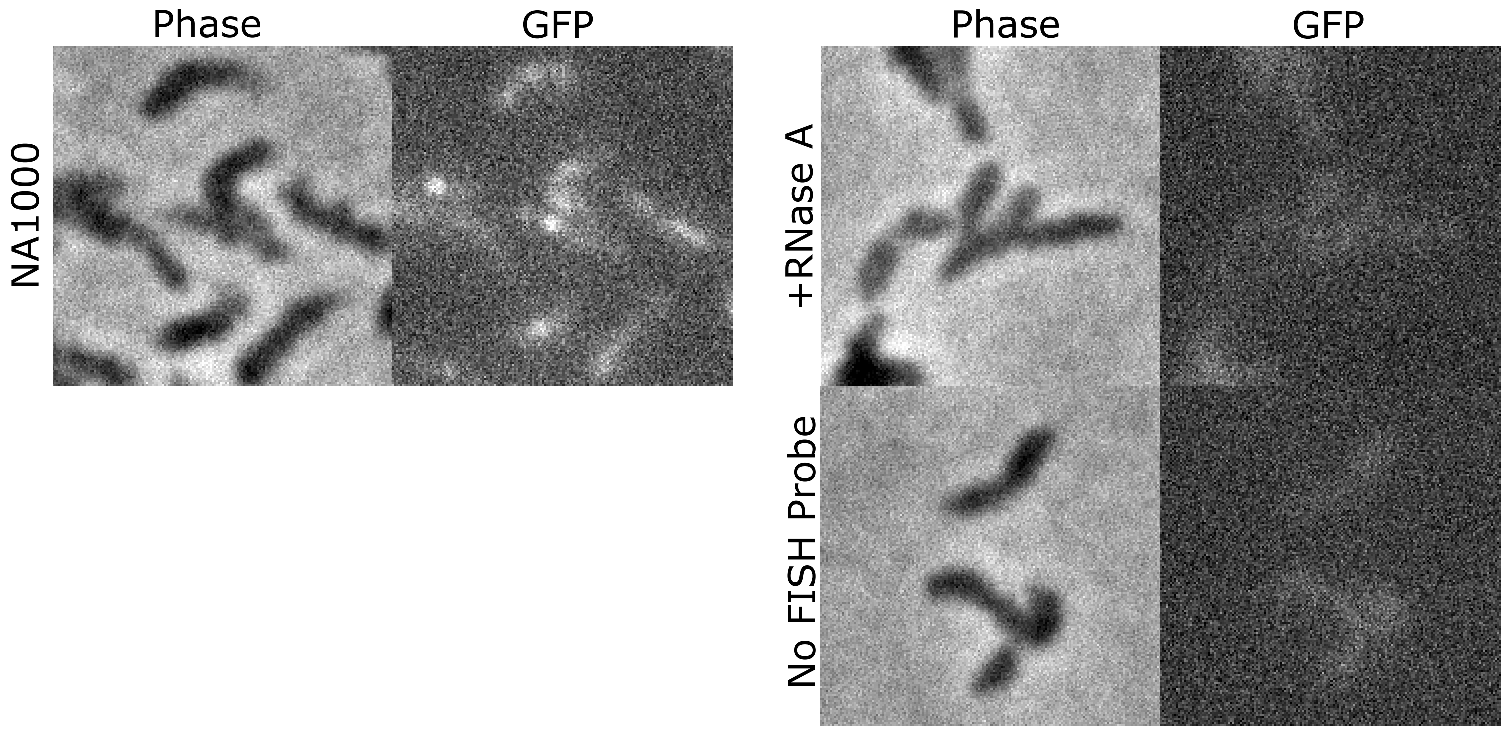
