## Supplementary material for "BR-bodies provide selectively permeable condensates that stimulate mRNA decay and prevent release of decay intermediates": Fig S3

A.

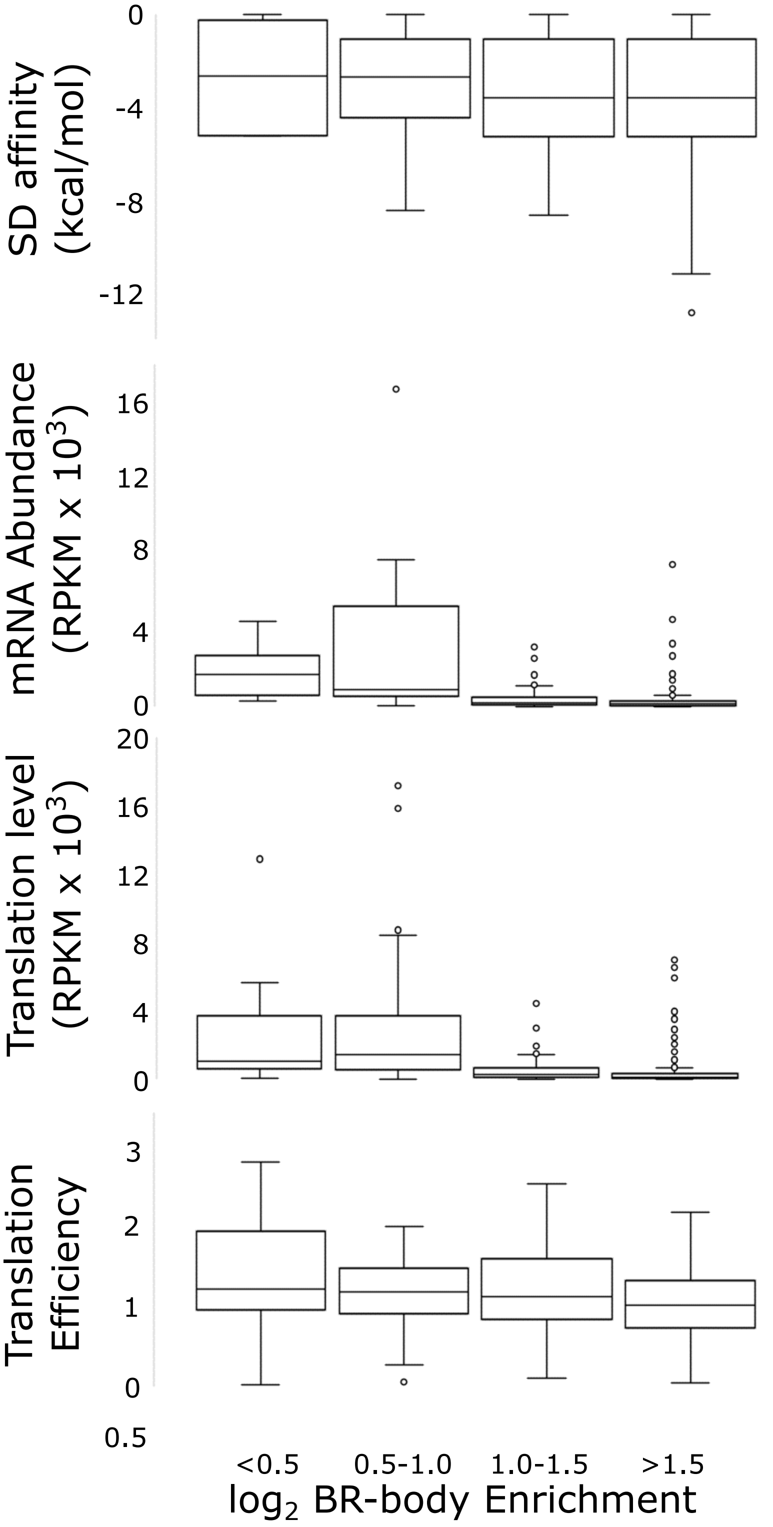


B.

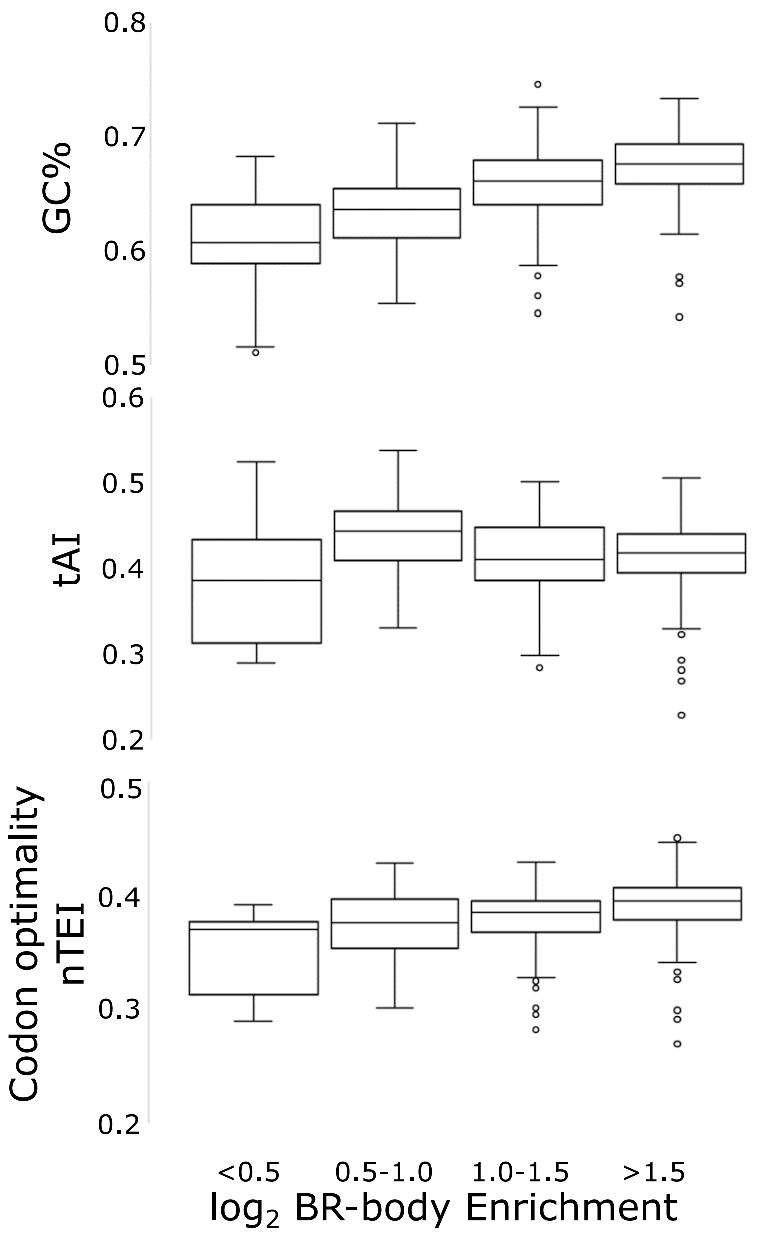


C.

| | BR-enrichment | mRNA half-life (min) | TE | mRNA abundance (RPKM) | mRNA Length (nt) | Translation level (RPKM) | GC% | TAI | 5' UTR length (nt) | nTE | SD calc ΔG (kcal/mol) |
|---|---|---|---|---|---|---|---|---|---|---|---|
| **BR-enrichment** | 1.00 | -0.15 | -0.16 | -0.51 | 0.41 | -0.40 | 0.47 | -0.01 | 0.02 | 0.30 | -0.04 |
| **mRNA half-life (min)** | -0.15 | 1.00 | 0.23 | 0.15 | -0.04 | 0.20 | -0.13 | 0.05 | 0.07 | -0.08 | -0.17 |
| **TE** | -0.16 | 0.23 | 1.00 | 0.11 | -0.03 | 0.24 | -0.10 | 0.19 | -0.01 | 0.05 | 0.01 |
| **mRNA abundance (RPKM)** | -0.51 | 0.15 | 0.11 | 1.00 | -0.23 | 0.87 | -0.31 | 0.25 | 0.06 | -0.12 | 0.03 |
| **mRNA Length (nt)** | 0.41 | -0.04 | -0.03 | -0.23 | 1.00 | -0.14 | 0.19 | 0.15 | 0.20 | 0.20 | -0.03 |
| **Translation level (RPKM)** | -0.40 | 0.20 | 0.24 | 0.87 | -0.14 | 1.00 | -0.18 | 0.25 | 0.10 | -0.07 | -0.02 |
| **GC%** | 0.47 | -0.13 | -0.10 | -0.31 | 0.19 | -0.18 | 1.00 | 0.24 | -0.02 | 0.57 | 0.06 |
| **TAI** | -0.01 | 0.05 | 0.19 | 0.25 | 0.15 | 0.25 | 0.24 | 1.00 | 0.16 | 0.51 | 0.01 |
| **5' UTR length (nt)** | 0.02 | 0.07 | -0.01 | 0.06 | 0.20 | 0.10 | -0.02 | 0.16 | 1.00 | -0.01 | -0.11 |
| **nTE** | 0.30 | -0.08 | 0.05 | -0.12 | 0.20 | -0.07 | 0.57 | 0.51 | -0.01 | 1.00 | 0.10 |
| **SD calc ΔG (kcal/mol)** | -0.04 | -0.17 | 0.01 | 0.03 | -0.03 | -0.02 | 0.06 | 0.01 | -0.11 | 0.10 | 1.00 |
